# Oncogenic RAS-Pathway Activation Drives Oncofetal Reprogramming and Creates Therapeutic Vulnerabilities in Juvenile Myelomonocytic Leukemia

**DOI:** 10.1101/2023.10.27.563754

**Authors:** Mark Hartmann, Maximilian Schönung, Jovana Rajak, Valentin Maurer, Ling Hai, Katharina Bauer, Mariam Hakobyan, Sina Staeble, Jens Langstein, Laura Jardine, Roland Rölz, Sheila Bohler, Eleonora Khabirova, Abdul-Habib Maag, Dominik Vonficht, Dirk Lebrecht, Katrin M. Bernt, Kai Tan, Changya Chen, Fatemeh Alikarami, Tobias Boch, Viktoria Flore, Pavlo Lutsik, Michael D. Milsom, Simon Raffel, Christian Buske, Simon Haas, Muzlifah Haniffa, Jan-Philipp Mallm, Sam Behjati, Marc-Jan Bonder, Stefan Fröhling, Charlotte M. Niemeyer, Joschka Hey, Christian Flotho, Christoph Plass, Miriam Erlacher, Matthias Schlesner, Daniel B. Lipka

## Abstract

Aberrant fetal gene expression facilitates tumor-specific cellular plasticity by hijacking molecular programs of embryogenesis^1^. Persistent fetal gene signatures in childhood malignancies are typically explained by their prenatal origins^2–6^. In contrast, reactivation of fetal gene expression is considered a consequence of oncofetal reprogramming (OFR) in adult malignancies and is associated with aggressive disease^7–10^. To date, OFR has not been described in the context of childhood malignancies. Here, we performed a comprehensive multi-layered molecular characterization of juvenile myelomonocytic leukemia (JMML) and identified OFR as a hallmark of aggressive JMML. We observed that hematopoietic stem cells (HSCs) aberrantly express mixed developmental programs in JMML. Expression of fetal gene signatures combined with a postnatal epigenetic landscape suggested OFR, which was validated in a JMML mouse model, demonstrating that postnatal activation of RAS signaling is sufficient to induce fetal gene signatures. Integrative analysis identified the fetal HSC maturation marker CD52 as a novel therapeutic target for aggressive JMML. Anti-CD52 treatment depleted human JMML HSCs and disrupted disease propagation *in vivo*. In summary, this study implicates OFR, defined as postnatal acquisition of fetal transcription signatures, in the pathobiology of a childhood malignancy. We provide evidence for the direct involvement of oncogenic RAS signaling in OFR. Finally, we demonstrate how OFR can be leveraged for the development of novel treatment strategies.

**Highlights:**

▪ Epigenomic and transcriptomic landscape of juvenile myelomonocytic leukemia (JMML) in the context of hematopoietic development.
▪ The presence of fetal transcription signatures in childhood malignancies is not indicative of a developmental maturation block.
▪ High-risk JMML is characterized by oncofetal reprogramming of postnatal hematopoietic stem cells (HSCs).
▪ RAS-pathway mutations induce fetal-like gene expression signatures in murine postnatal HSCs.
▪ The fetal maturation marker CD52 is a novel therapeutic target in high-risk JMML.

## Introduction

Fetal signatures in cancer have been described for a number of entities including pediatric and adult malignancies, although the cellular origins and molecular mechanisms are mostly obscure^9,10^. For childhood malignancies, the detection of driver mutations at the time of birth revealed that premalignant clones can arise prenatally leading to a maturation block with subsequent malignant transformation^11^. However, genetic evidence for a prenatal event cannot be obtained in all patients affected by childhood malignancies, suggesting that alternative molecular mechanisms may establish fetal signatures. In adult malignancies, the concept of ‘oncofetal reprogramming’ (OFR) describes the reactivation of fetal programs in cancer cells originating from postnatal cells-of-origin. OFR could explain the occurrence of fetal expression programs in childhood malignancies for which evidence of a prenatal origin is lacking. However, to date, OFR has not been described in the context of childhood cancer^1,11^.

Juvenile myelomonocytic leukemia (JMML) is a myeloproliferative/myelodysplastic neoplasm of early childhood and clinically a highly heterogeneous disease that is defined by the presence of mutations affecting the RAS-signaling pathway^12^. Epigenetic disease subgroups, known as epitypes, have been identified as the only solitary significant predictor of overall survival in JMML^13–16^. In addition to global DNA hypermethylation, the upregulation of fetal hemoglobin and the fetal hematopoietic stem cell (HSC) marker *LIN28B* have been associated with high-risk disease^17–19^. Correspondingly, in a subset of JMML patients oncogenic driver mutations have been found in neonatal blood spots, suggesting a prenatal origin of the disease^20^. However, JMML driver mutations can only be detected in a minority of high-risk patients at birth, which may indicate a postnatal origin of the disease in these patients^21^. Such indications of a postnatal disease origin in some cases together with the simple genetic landscape and the profound epigenetic remodeling qualifies JMML as a suitable model to study cancer ontogeny.

In this study, we used multi-omics profiling to identify disease-specific transcriptomic and epigenomic aberrations associated with high-risk JMML. We observed an unanticipated discrepancy between prenatal transcription and postnatal methylation states, suggesting OFR as a possible explanation. Using a JMML mouse model driven by HSC-specific activation of a *Ptpn11^E76K^* mutation revealed that oncogenic RAS pathway activation results in OFR. Furthermore, we demonstrate that OFR drives disease-specific expression programs, resulting in aberrantly upregulated markers in HSCs of high-risk JMML patients. Preclinical data testing one of these aberrantly expressed markers in a xenotransplantation model suggested that CD52 may serve as an efficient therapeutic target in high-risk JMML.

## Results

### JMML-related aberrations affect the entire hematopoietic system including hematopoietic stem cells

The clinical heterogeneity found across JMML patients has mostly been resolved at the molecular level using DNA methylation analysis which led to the identification of so-called JMML epitypes^13–15^. However, the molecular mechanisms driving the establishment of JMML epitypes remain elusive. To systematically characterize the degree of molecular heterogeneity in JMML, we adopted a multi-modal analysis strategy to integrate epigenomic, transcriptomic, and surface marker expression data obtained from patient samples representing all three JMML epitypes: namely low (LM); intermediate (IM); and high methylation (HM) JMML (Fig. 1a, Extended Data Fig. 1, Supplementary Table 1)^16^. Using scRNA-seq of mononuclear cells (MNC) isolated from JMML patients and healthy subjects of different age groups (Extended Data Fig. 2, Supplementary Table 2), we confirmed that all hematopoietic lineages and major cell types were present in the hematopoietic systems of JMML patients, although cell type frequencies were heterogenous across patients (Fig. 1b-d, Extended Data Figs. 3 and 4).

**Fig. 1:**
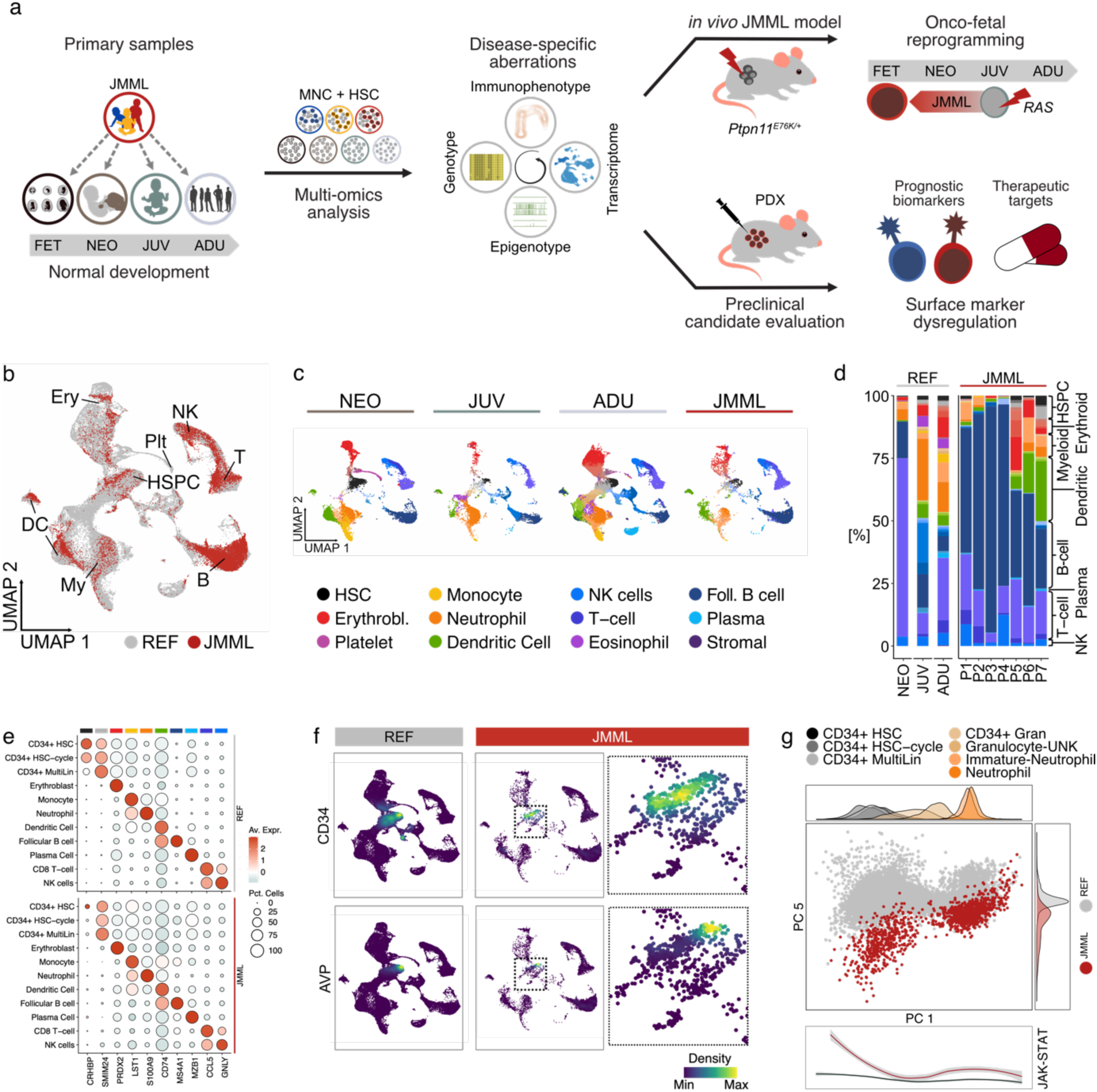
JMML-related aberrations affect the entire hematopoietic system including hematopoietic stem cells. **a**, Conceptual and experimental overview. Cellular and molecular aberrations in JMML are characterized in the context of healthy hematopoietic development using a multi-omics approach to identify disease-specific aberrations that can be used for diagnostic and therapeutic purposes. (FET: fetal; NEO: neonatal; JUV: juvenile; ADU: adult; MNC: mononuclear cells; HSC: hematopoietic stem cells; PDX: patient-derived xenograft). **b-g**, Single-cell RNA-sequencing (scRNA-seq) of mononuclear cells (MNCs) isolated from JMML patient samples and healthy postnatal references. Cell types were annotated using adult human cell atlas data as a reference. **b**, UMAP of scRNA-seq data colored by disease status (grey: healthy references; red: JMML). ‘REF’ includes data from neonatal, juvenile and adult samples. **c**, UMAPs of scRNA-seq data split by dataset and colored by cell type. Major cell types are indicated in the legend, detailed cell type annotation is included in **Extended Data** Fig. 3). Erythrobl.: erythroblast; Foll. B cell: follicular B cell; Plasma: plasma cell. **d**, Stacked bar plot summarizing the cell type frequencies across the scRNA-seq datasets. P1 – P7: JMML samples. **e**, Dot plot summarizing gene expression of characteristic lineage markers across major hematopoietic cell types in healthy references (REF) and JMML samples. **f**, Density UMAPs depicting the expression of the hematopoietic stem and progenitor cell markers *AVP* and *CD34*. **g**, Principal component analysis (PCA) of the neutrophil differentiation trajectory. Correlation analysis identified PC 1 and PC 5 as representative of neutrophil differentiation (grey to orange) and disease-specific features (red), respectively. Cells from healthy references: grey; cells from JMML patients: red. Density plots depict the cell density along the axes stratified by cell type (PC 1) or by disease status (PC 5). Bottom panel describes JAK-STAT signaling pathway activity across neutrophil pseudotime for JMML (red) and references (black).

However, pronounced differences were observed between JMML samples and healthy controls regarding the frequency of cells expressing typical age-matched hematopoietic stem and progenitor cell (HSPC) marker genes (Fig. 1e,f, Extended Data Figs. 3 and 4). These aberrant expression profiles were also observed in a systematic trajectory analysis and affected all major lineages, including hematopoietic stem cells (HSC; Fig. 1g, Extended Data Fig. 5). All trajectories showed enrichment of inflammatory signaling pathways with a peak in HSCs (Fig. 1g, Extended Data Fig. 5d). Together, this data suggested that the HSC compartment is massively altered in JMML.

### Epitypes are conserved in hematopoietic stem cells of JMML patients

To systematically characterize early molecular evolution of JMML epitypes, we investigated the molecular landscape of the HSPC compartment of JMML patients by performing a multi-modal analysis comprising immunophenotypes, epigenotypes, and transcriptomes (Fig. 2a, Extended Data Fig. 6). Lineage-negative (Lin^-^) cells of JMML patients revealed cell surface expression patterns for CD34 and CD38 that were different in frequencies but comparable in global structure to those observed in healthy references (Fig. 2b, top panel). In contrast, Lin^-^CD34^+^CD38^-/lo^ HSPCs exhibited highly heterogeneous immunophenotypes with respect to CD45RA and CD90 expression (Fig. 2b, bottom panel). This included the appearance of a recently described aberrant Lin^-^CD34^+^CD38^-/lo^CD45RA^+^CD90^+^ HSPC population^22,23^. The abundance of these immunophenotypically aberrant CD45RA^+^CD90^+^ HSPCs was distinct across JMML epitypes, with hypermethylated (HM) JMML showing the highest frequency of CD45RA^+^CD90^+^ HSPCs in an independent validation cohort (n=20 pts; Fig. 2c, Supplementary Table 3). This observation suggested that epitype-related, disease-specific aberrations of gene regulatory programs might already be present in the HSPC compartment of JMML patients. To test this possibility, we generated ultra-low input whole-genome bisulfite sequencing (WGBS) data from different HSPC subpopulations, i.e. the CD45RA/CD90 quadrants of Lin^-^CD34^+^CD38^-/lo^ cells (Fig. 2b, bottom panel, Supplementary Table 4). DNA methylation-based cell type classification^24^ determined all CD45RA/CD90 subpopulations isolated from JMML patients as HSCs, independent of the epitype, the donor, or the immunophenotypic CD45RA/CD90 quadrant, confirming aberrant molecular programs in the JMML HSC compartment (Extended Data Fig. 7a). In line with this observation, unsupervised analysis of JMML HSC methylomes precisely recapitulated the corresponding epitypes that had been determined by methylation array analysis of bulk MNCs (Fig. 2d, Extended Data Figs. 1 and 7b-d). In conclusion, the conservation of epitypes between HSCs and differentiated MNCs from JMML patients reveals DNA methylation as a differentiation-independent and early disease-specific aberration, which confirms JMML as a disease originating from HSCs.

**Fig. 2:**
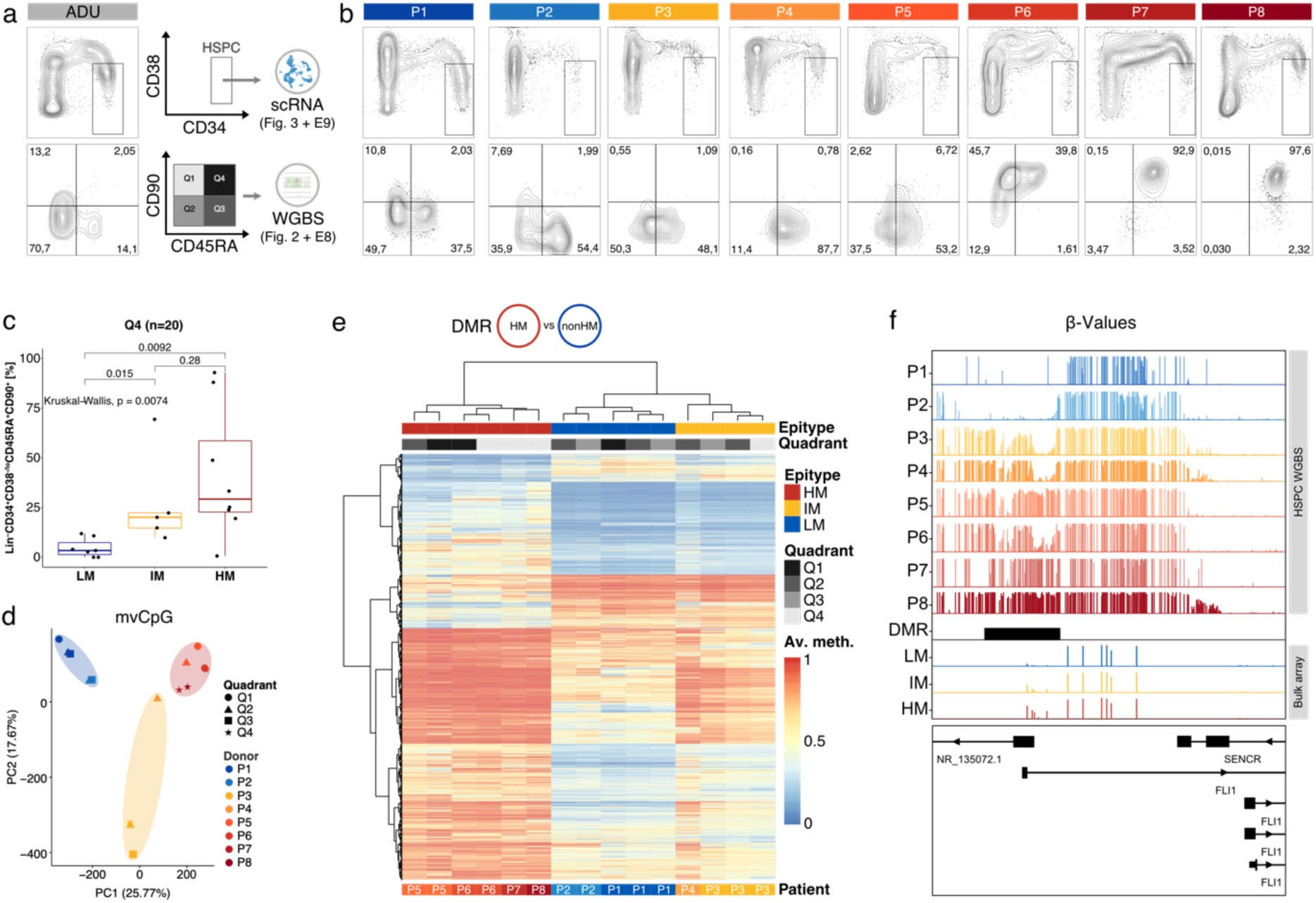
Epitypes are conserved in hematopoietic stem cells of JMML patients. **a**, Left: FACS contour plots from healthy adult bone marrow (n=1; ADU). Right: schematic representing the sorting strategy of HSPCs for single-cell RNA-sequencing (top; Lin^-^CD34^+^CD38^-/lo^) and ultra-low input whole-genome bisulfite sequencing (Lin^-^CD34^+^CD38^-/lo^CD45RA/CD90 quadrants). scRNA: single-cell RNA-sequencing; WGBS: ultra-low input whole-genome bisulfite sequencing. **b**, FACS contour plots depicting expression of CD38 (y-axis) over CD34 (x-axis) in the Lin^-^ (top) and expression of CD90 (y-axis) over CD45RA (x-axis) in the Lin^-^CD34^+^CD38^-/lo^ (bottom) compartments of JMML patients (n=8; P1 – P8). Colors refer to the semi-quantitative genome-wide methylation states in each patient (Extended Data Fig. 1). **c**, Quantification of aberrant Lin^-^ CD34^+^CD38^-/lo^CD45RA^+^CD90^+^ HSCs in a validation cohort (n=29 JMML patients) stratified by JMML epitype. **d-f**, µWGBS of sorted JMML HSPCs. **d**, Principal component analysis (PCA) of the 20,000 most-variable CpGs identified across JMML HSPCs. **e**, Heatmap depicting methylation status of 13,832 DMRs identified between HM and non-HM (i.e., “intermediate methylation” [IM] and “low methylation” [LM]) patients. Hierarchical clustering using Manhattan distance and complete linkage. **f**, Genome browser tracks of WGBS and bulk methylation array data. Tracks P1 – P8 depict WGBS data from JMML HSPCs. LM, IM, and HM refer to bulk methylation array data aggregated from 147 JMML patients representing all epitypes (LM=62, IM=45, HM=40). Depicted is the genomic region (hg19, chr11:128,553,548-128,564,856) containing a DMR overlapping an alternative transcription start site of *FLI1*.

Next, we aimed to identify DNA methylation changes in JMML HSCs that are characteristic of the HM epitype, as this epitype is associated with an aggressive disease course^13,16^. Differentially methylated regions (DMRs) detected between HSCs from HM and non-HM (LM & IM) patients confirmed widespread CpG hypermethylation, which particularly affected bivalent enhancers and polycomb-repressed regions associated with developmental processes (Fig. 2e, Extended Data Fig. 7b and e-l, Supplementary Table 5).

For example, DMRs were annotated to loci of hematopoietic and developmental regulators such as *FLI1*, *RUNX1*, and several homeobox gene classes (Fig. 2f, Extended Data Fig. 7m,n). Furthermore, DMRs were enriched for transcription factor (TF) binding motifs such as PU.1, HOXC9, as well as various GATA and ETS family members, suggesting an impact of epitypes on specific gene regulatory programs involved in hematopoietic maturation and differentiation (Extended Data Fig. 7o). In conclusion, our DNA methylome data demonstrated that JMML epitypes are present in HSCs from JMML patients, and that the observed epigenetic aberrations might govern epitype-specific gene regulation, including the engagement of developmental programs.

### Hematopoietic stem cells exhibit epitype-specific gene expression programs in JMML patients

To investigate whether the epitype-specific DNA methylation patterns translate into distinct gene expression programs in HSCs of JMML patients, we performed scRNA-seq on Lin^-^CD34^+^CD38^-/lo^ HSPCs isolated from the same 8 JMML patients (P1 – P8; Fig. 2a and Extended Data Fig. 8a, Supplementary Table 2). Focusing on transcriptionally defined HSCs, we found significant transcriptional priming towards the myeloid lineage (p = 0.0023, LMM and ANOVA) in patients with IM and HM epitypes as compared to the LM epitype, and this increase in transcriptional myeloid priming occurred at the expense of priming towards the megakaryocytic-erythroid lineages (Extended Data Fig. 8b, Supplementary Table 6). These differences were paralleled by an increase in expression of leukemia-associated signatures from LM- to HM-HSCs, suggesting that aberrant DNA methylation might impact on HSC function in JMML in an epitype-specific manner (Extended Data Fig. 8c, Supplementary Table 6). Differential gene expression analysis between HSCs from HM and non-HM patients revealed a number of highly dysregulated genes, which are critical for hematopoiesis, including HSC markers such as *AVP*, *CRHBP*, *CD164*, and *CD34*, differentiation and activation markers such as *LGALS1*, *DNTT*, *IGLL1*, *CD96*, *CD69*, and *CD52*, as well as developmental factors such as *HMGA2* and *HBG2* (Extended Data Fig. 8d, Supplementary Table 7). Functional enrichment analyses confirmed immune activation and inflammation programs in JMML HSCs, which has been described as a feature of prenatal development^25^ (Extended Data Fig. 8e,f, Supplementary Table 8-11). Taken together, the upregulation of developmental genes in HM-JMML HSCs was in line with the altered DNA methylation seen in regions that are epigenetically regulated during development (Extended Data Fig. 7h-o). This indicated an epitype-specific functional role for developmental factors in JMML pathogenesis and raised the question whether developmental origins vary across JMML epitypes.

### Transcriptional mosaicism of developmental programs in JMML HSCs

A recent report described the detection of JMML driver mutations at birth in 75% (12 out of 16) of individuals who later developed JMML (Extended Data Fig. 8g)^21^. Together with our observation that JMML epitypes are associated with developmental features in HSCs, this suggested a prenatal origin of the disease in at least a subset of patients. To systematically examine developmental phenotypes of JMML HSCs and to compare these to healthy development, we compiled a scRNA-seq reference map of hematopoietic maturation, including 116 different reference cell types from healthy donors spanning four developmental stages: fetal (FET), neonatal (NEO), juvenile (JUV), and adult (ADU; Fig. 3a, Supplementary Table 2). Next, we applied “cell type similarity inference”^5^ to independently predict the average similarity of JMML HSCs per patient to each of the reference cell types (Extended Data Fig. 8h). While HSCs from all JMML samples showed high similarity to healthy neonatal HSCs, LM-JMML HSCs also exhibited high similarity to healthy postnatal (i.e., juvenile and adult) HSCs. In contrast, the IM- and HM-JMML stem cells showed high transcriptional similarity to healthy fetal HSCs (Fig. 3b). Such subgroup-specific global expression patterns were confirmed by the expression of known marker genes for HSCs from different developmental stages. Postnatal HSC markers (*CRHBP*, *AVP*) were found to be upregulated in LM-JMML and healthy postnatal HSCs (Fig. 3c, Extended Data Fig. 8i). Conversely, fetal HSC markers, such as *HMGA2* and *MECOM*, were upregulated in HM-JMML and healthy fetal HSCs (Fig. 3d, Extended Data Fig. 8i), which implied either preservation or reactivation of fetal gene expression programs in HM-JMML. In this context, it is noteworthy that JMML HSCs revealed different levels of similarity (>0.5) but virtually no dissimilarities (<0.5) to any of the reference HSC populations across hematopoietic development (Fig. 3b, Extended Data Fig. 8h). This suggested a transcriptional mosaicism of developmental signatures rather than an unambiguous cell type-specific transcriptional profile for any of the stages. To consolidate this observation, we generated a list of the top 50 stage-specific marker genes of normal HSC development and analyzed the expression of these signature genes in JMML HSCs (Fig. 3e, Supplementary Table 12-13). This revealed a high degree of subgroup- and patient-specific heterogeneity, including aberrant regulation of a range of genes relative to normal HSCs from all of the four developmental stages. This further suggested that JMML is characterized by variegated expression of developmental signatures, including components of fetal gene expression programs, most prominently in HM-JMML.

**Fig. 3:**
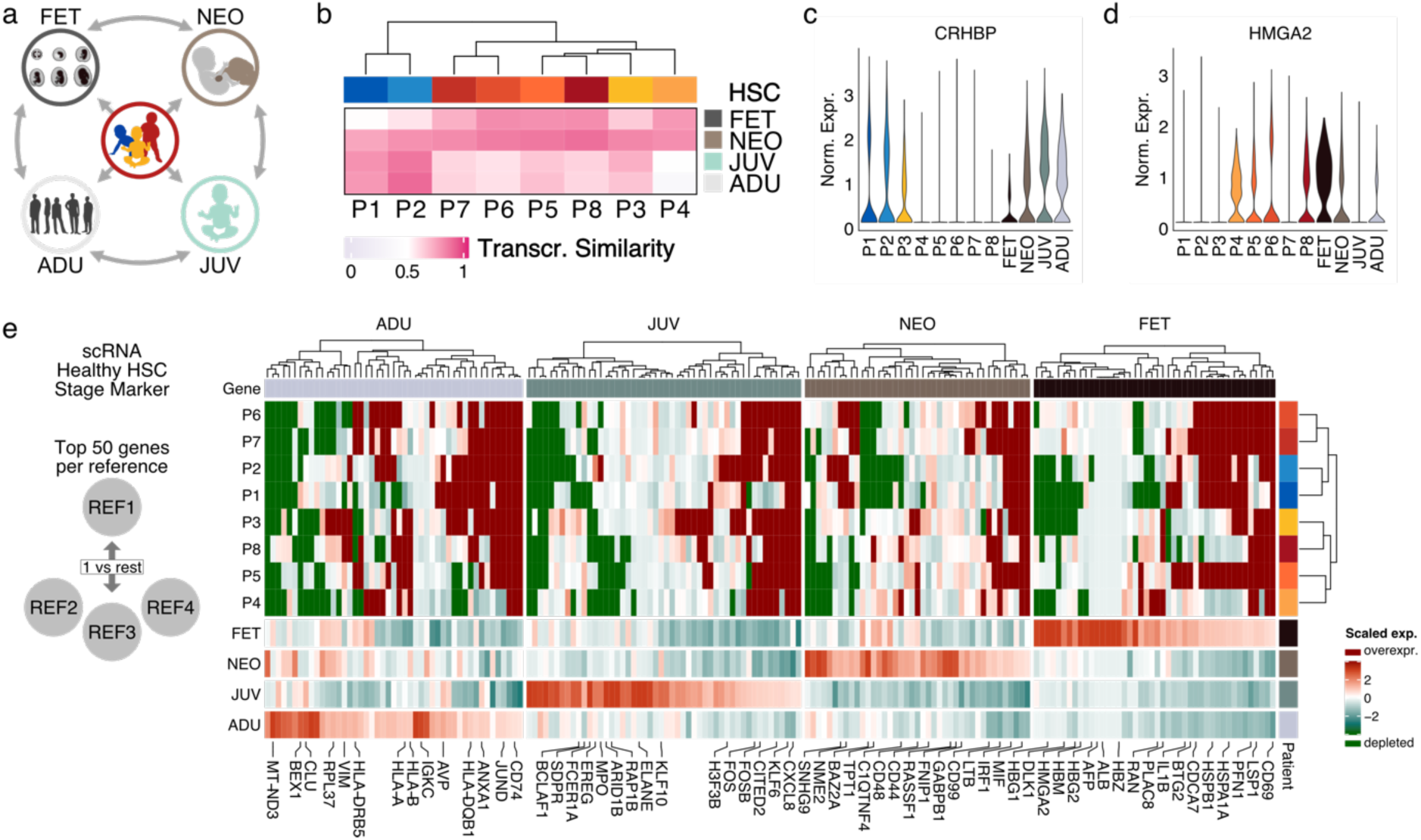
JMML HSCs are characterized by transcriptional mosaicism of developmental programs. **a**, Schematic displaying the systematic comparison of JMML HSCs to healthy HSCs across different developmental stages including fetal liver (FET), cord blood (NEO), juvenile (JUV), and adult (ADU) bone marrow. **b-e**, Transcriptional profiling using scRNA-seq of JMML HSCs in the context of healthy HSCs representing different developmental stages. HSCs were defined uniformly based using automated cell type annotation (see “Materials and Methods” section). **b**, Cell type similarity inference using logistic regression analysis. JMML HSCs were compared at the single-cell level to healthy HSCs isolated at different developmental stages. Depicted are averaged and probability-converted predicted logits. Values >0.5 (pink) indicate similarity whereas values <0.5 (grey) indicate dissimilarity to the references (rows). **c-d**, Violin plots showing the expression profiles of the HSC marker genes *CRHBP* and *HMGA2* in healthy and malignant HSCs. *CRHBP* is a marker for adult HSCs and *HMGA2* is a marker for fetal HSCs. **e**, Heatmap depicting expression profiles of stage-specific signature genes for healthy HSC development. For each developmental stage, the top 50 DEGs compared to all other stages were identified. Only genes, that were found to be differentially expressed in a single reference were kept. Scaling of normalized expression was done on a per gene basis across healthy HSC references to reflect the physiologic expression range. All values above the maximum or below the minimum are flagged with dark red or dark green, respectively, representing aberrantly up- or downregulated signature genes in JMML.

### DNA methylome analysis reveals postnatal maturation of hematopoietic stem cells across all JMML epitypes

The observed transcriptional mosaicism in JMML HSCs was inconclusive with respect to the developmental origins of JMML. Since it has previously been demonstrated that DNA methylomes retain elements of the tissues and cell types of origin^26^, we sought to investigate the DNA methylomes of JMML HSCs in the context of the developing hematopoietic system. We generated a DNA methylome reference map of normal HSC maturation representing the same developmental stages as used for the scRNA-seq analysis, i.e. fetal, neonatal, juvenile, and adult HSCs (Fig. 4a, Supplementary Table 4). Unsupervised analysis of the 10,000 most variable CpGs across normal and JMML HSCs revealed pronounced genome-wide differences between HM-JMML and healthy references from all developmental stages, whereas LM-JMML HSCs clustered with the postnatal references (Extended Data Fig. 9a). To more precisely determine patterns associated with HSC maturation, we identified genomic regions with dynamic DNA methylation in healthy references and found 1,179 regions (441 with methylation gain, 738 with methylation loss) that revealed continuous gain or loss of DNA methylation from fetal to adult HSCs (Fig. 4b, Supplementary Table 14). Using these maturation dynamic regions for phylogenetic analysis recapitulated the trajectory from fetal to adult HSCs during normal maturation (Extended Data Fig. 9b). Projecting the JMML HSC methylomes onto this maturation phylogeny revealed postnatal epigenetic patterns for all JMML HSCs analyzed here, regardless of their epitype, indicating that at the time of diagnosis all JMML HSCs had acquired a postnatal DNA methylome (Fig. 4c, Extended Data Fig. 9c). This finding was further corroborated when looking at so called ‘epigenetic scars’, which are genomic regions that change their methylation status significantly in one particular developmental window but remain stable thereafter (Fig. 4d and Extended Data Fig. 9d, Supplementary Table 15-16). These analyses showed that HSCs from all JMML patients clustered together with postnatal HSCs when looking at the transitions from fetal to postnatal or from neonatal to post-neonatal HSCs, respectively, further indicating that HSCs from all patients examined had passed the neonatal stage regardless of their epitype (Fig. 4e and Extended Data Fig. 9e). In conclusion, our epigenetic data show postnatal epigenomes in HSCs from all 8 JMML patients investigated, which is in contrast to the fetal-like expression signatures observed in non-LM-JMML. These seemingly contradictory findings could theoretically be explained by two different models: (I.) Prenatal acquisition of a RAS-pathway mutation could lead to altered HSC maturation resulting in partial preservation of fetal transcription signatures (fetal origin hypothesis; Fig. 4f, left panel). (II.) Alternatively, acquisition of a RAS-pathway mutation in a postnatal HSC may lead to transformation and induction of aberrant transcription programs which result in reprogramming and reactivation of fetal-like molecular signatures (OFR hypothesis; Fig. 4f, right panel).

**Fig. 4:**
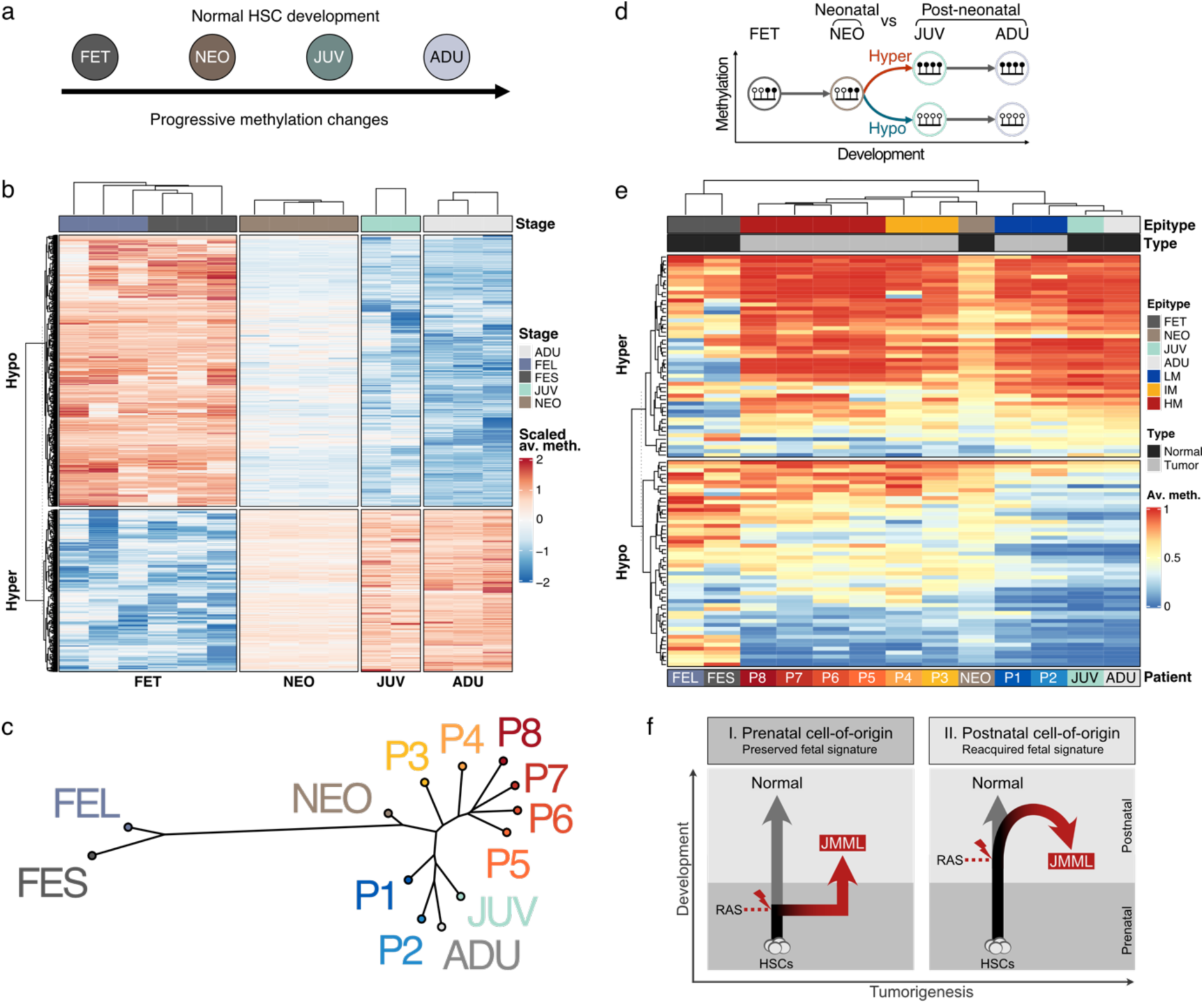
DNA methylome analysis reveals a postnatal maturation stage in JMML HSCs across all epitypes. DNA methylome analysis of JMML HSCs in the context of healthy hematopoietic development using ultra-low input WGBS. Fetal HSCs (*FET*; fetal spleen HSCs [n=3], fetal liver HSCs [n=3]), cord blood HSCs (*NEO*; n=4), juvenile HSCs (*JUV*; n=2), and adult HSCs (*ADU*; n=3). **a**, Schematic summarizing the normal maturation trajectory of healthy HSCs, which was used to determine maturation-associated methylation changes. **b**, Heatmap showing the hierarchical clustering of all methylation-dynamic regions (n=1,179) identified in HSCs across normal development. Plotted are scaled average DNA methylation values. **c**, Phylogenetic tree analysis based on all methylation-dynamic regions. Methylome data of JMML HSCs are projected onto this phylogenetic tree based on Manhattan distances. Replicates are aggregated per patient or healthy developmental stage. **d-e**, ‘Epigenetic scars’ analysis from the transition between neonatal and postneonatal stages. **d**, Epigenetic scars are defined as DNA methylation changes that occur in a given developmental window but do not change thereafter. **e**, Heatmap showing methylation beta values of epigenetic scars from the transition of neonatal to post-neonatal HSCs. Hypo: loss of DNA methylation; hyper: gain of DNA methylation. **f**, Schematic illustrating the two hypothetical scenarios explaining the expression of fetal-like signatures in JMML HSCs: I. Transformation of a prenatal cell-of-origin reprograms HSCs leading to a partial preservation of fetal HSC programs in JMML patients (left panel). II. Alternatively, transformation of a postnatal HSC partially reactivates fetal gene expression patterns, resulting in oncofetal reprogramming (OFR) (right panel).

### RAS-pathway activation in postnatal HSCs results in oncofetal reprogramming in a JMML mouse model

To experimentally test whether postnatal acquisition of a JMML driver mutation is able to instruct the reactivation of fetal-like expression signatures (OFR hypothesis), we established an inducible JMML mouse model which is driven by the activation of a *Ptpn11^E76K^* mutation in HSCs upon tamoxifen injection (Fig. 5a, Extended Data Fig. 10a). Postnatal induction of the *Ptpn11^E76K^*mutation in adolescent mice (6-12 weeks of age) resulted in rapid development of a JMML-like disease, characterized by leukocytosis, thrombocytopenia, reduced hemoglobin levels, pronounced splenomegaly, excess production of myeloid cells, particularly monocytes, and a median survival of 24 weeks (Fig. 5b-f, Extended Data Fig. 10b-e). To analyze the molecular mechanisms underlying the JMML-like disease observed in *Ptpn11^E76K/+^* mice, we isolated LSK (Lin^-^Sca1^+^cKit^+^), LK (Lin^-^cKit^+^), and total hematopoietic cells from the bone marrow (BM) of *Ptpn11^E76K/+^* and *Ptpn11^+/+^* mice 6 weeks after tamoxifen induction and performed scRNA-seq (Supplementary Table 17). Data were integrated across genotypes and annotated based on the expression of established marker genes (Fig. 5g, Extended Data Fig. 10f). We found a significant myelomonocytic skewing and loss of immature HSPCs in *Ptpn11^E76K/+^* mice (Fig. 5h, Extended Data Fig. 10g,h), demonstrating that the postnatal induction of *Ptpn11^E76K/+^*in HSCs precisely recapitulated human JMML. The upregulation of monocytic gene expression programs was already observed at the level of HSCs in *Ptpn11^E76K/+^*mice, indicating increased transcriptional priming of *Ptpn11^E76K/+^*HSCs towards the myeloid lineage (Fig. 5i, Extended Data Fig. 10i-k, Supplementary Table 6). In addition, we found a significant enrichment of fetal gene signatures in HSCs from *Ptpn11^E76K/+^* mice as compared to wild-type HSCs (Fig. 5j,k, Supplementary Table 6). In line with this observation, *Ptpn11^E76K/+^*HSCs showed a significant upregulation of fetal HSC markers such as *Lgals1* and *Plac8* and concurrent downregulation of adult HSC marker genes (*H3f3b*, *Fos,* and *Procr;* Fig. 5l).

**Fig. 5:**
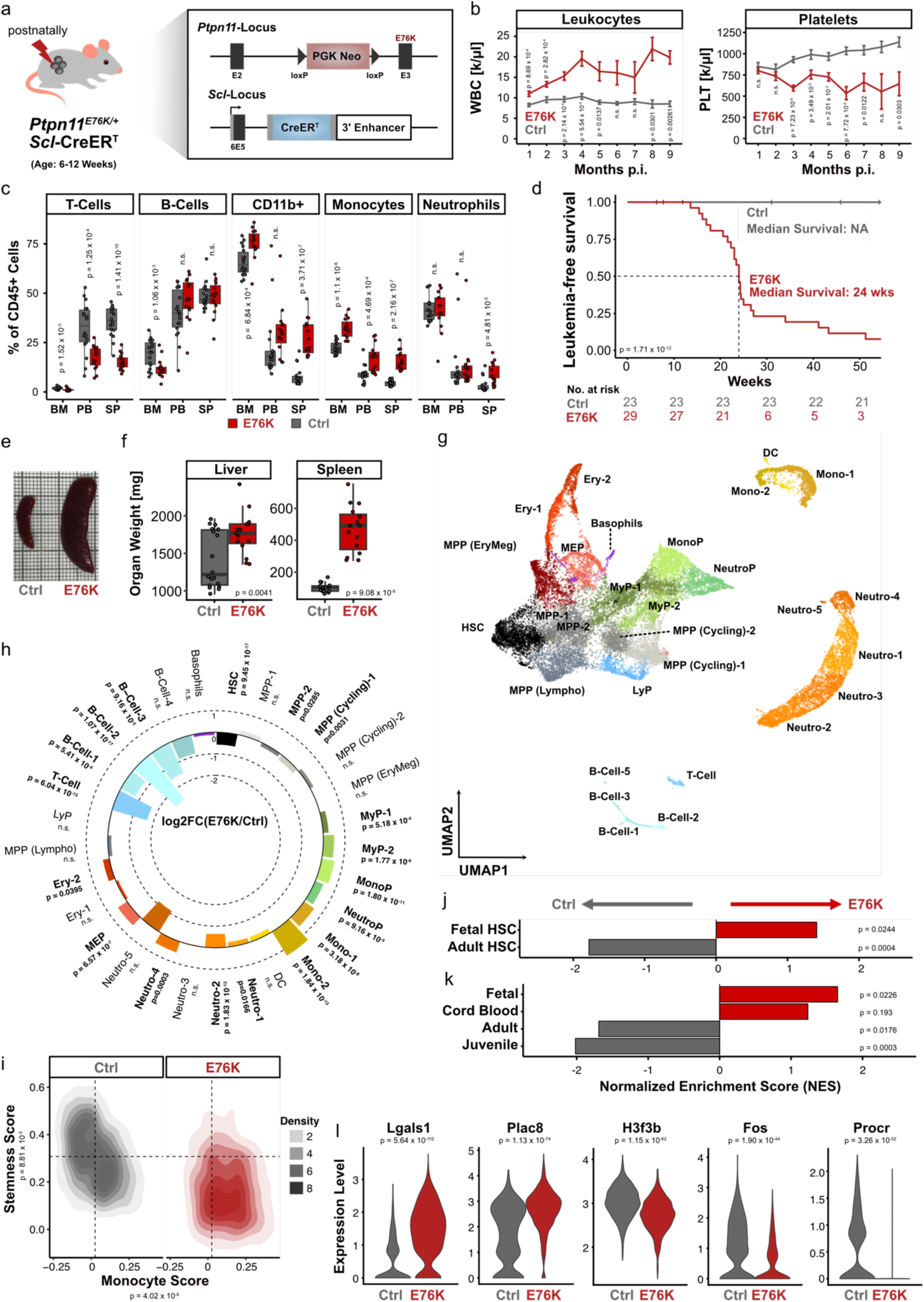
RAS-pathway activation in postnatal HSCs results in oncofetal reprogramming in a JMML mouse model. **a**, Establishment of a JMML mouse model by induction of a *Ptpn11*^E76K^ mutation in postnatal HSCs. Conditional *Ptpn11*^E76K/+^ mice were crossed to SclCreER^T^ mice to allow HSPC-specific activation of the mutation. *Ptpn11*^E76K/+^ Scl-CreER^T^ mice (E76K) and Ptpn11^+/+^ Scl-CreER^T^ controls (Ctrl) were injected (i.p.) with tamoxifen (TAM) at the age of 6-12 weeks. **b**, Leukocyte (WBC) and platelet (PLTs) counts are shown throughout 9 months post induction (p.i.). Mean and SEM are depicted. Statistical significance was calculated using student’s t-test. **c**, Differentiated hematopoietic cells were measured by flow cytometry in bone marrow (BM), spleen (SP) and peripheral blood (PB) of mice 18 weeks p.i. The box plots show the abundance of each cell type as percentage of total CD45^+^ cells. Whiskers are defined as 1.5 times interquartile range (IQR). Statistical significance was calculated using Student’s t-test. **d**, Kaplan-Meier plot depicting leukemia-free survival. Numbers of animals at risk are denoted under the respective timepoints. Median survival is indicated and significance of survival differences was calculated using log-rank test. **e-f**, Spleens and livers were isolated 18 weeks p.i. and organ weights were documented. Whiskers are defined as 1.5 times interquartile range (IQR). Statistical significance was calculated using Student’s t-test. **g**, Single-cell RNA-seq of E76K (n=3) and Ctrl (n=3) mice 18 weeks after TAM induction. The UMAP shows a 2D representation of 29 clusters identified and annotated to a hematopoietic cell type based on marker gene expression. **h**, Circular bar plot depicting the relative changes in cluster contribution of E76K compared to Ctrl cells. Significance was determined using Fisher’s exact test followed by Benjamini-Hochberg’s FDR correction. **i**, Density plot displaying transcriptional priming of HSCs towards the monocytic lineage (“Monocyte Score”) compared to expression of a “stemness” signature (“Stemness Score”). Significance was determined based on LMM and subsequent ANOVA. **j**, Enrichment analysis of mouse fetal and adult HSC signatures ^27^ in E76K and Ctrl HSCs. **k**, Enrichment analysis of human HSC signatures across different developmental stages (signature genes identified in this study, Fig. 3e) HSCs. **l**, Violin plots showing the expression of marker genes for different developmental stages of the hematopoietic system in HSCs. Significance was determined using Wilcoxon rank sum test.

In summary, our mouse model recapitulated phenotypic and molecular features of human JMML including transcriptional rewiring and myeloid lineage priming of HSCs. We demonstrated that expression of the *Ptpn11^E76K^* mutation in adolescent mice was sufficient to re-establish fetal-like gene expression programs in HSCs, hence providing proof-of-principle that postnatal acquisition of a RAS-pathway mutation can lead to oncofetal reprogramming.

### Integrative analysis identifies novel prognostic biomarkers and therapeutic targets for high-risk JMML

The mouse model demonstrated that HSC-specific activation of *Ptpn11^E76K^*^7^*^/+^* induces a JMML-like disease. This provided further evidence that HSCs are the likely cell-of-origin of JMML and suggested that effective therapies should target aberrant molecular features of JMML HSCs. Following the experimental proof of OFR in JMML, we asked the question whether disease-specific aberrations affect developmental factors that could be exploited for the treatment of JMML. To identify and prioritize such aberrations, we defined the following selection criteria: (a) DNA methylation-associated gene expression in JMML HSCs; (b) aberrant upregulation of gene expression in JMML HSCs relative to healthy reference HSCs across developmental stages; (c) proposed role in HSC development; (d) availability of drugs, which can be used for preclinical testing (Fig. 6a). To identify DNA methylation-associated genes in HSCs across JMML epitypes, we determined genes for which differential expression could be explained by differential DNA methylation. Comparing HM with non-HM JMML, we identified 346 DMRs that are associated with 155 DEGs (out of 255 HM vs non-HM DEGs overall) (Extended Data Fig. 11a, Supplementary Table 18-19), which confirmed the central role of DNA methylation in disease-specific gene regulation. For functional validation, we defined high-confidence candidate genes based on high correlation (>0.9) of DNA methylation and gene expression (Extended Data Fig. 11a, Fig. 6b). Out of 56 high-confidence candidates, 23% (13 genes) encoded for cell surface proteins (Fig. 6b, Extended Data Fig. 11b), which is significantly more than the expected proteome-wide frequency^28^. Comparison of expression levels revealed that all 13 candidates were consistently upregulated in JMML HSCs relative to almost all healthy reference HSCs. Moreover, these surface proteins showed epitype-specific expression patterns, suggesting that they could act as both disease- and epitype-specific biomarkers (Extended Data Fig. 11c).

**Fig. 6:**
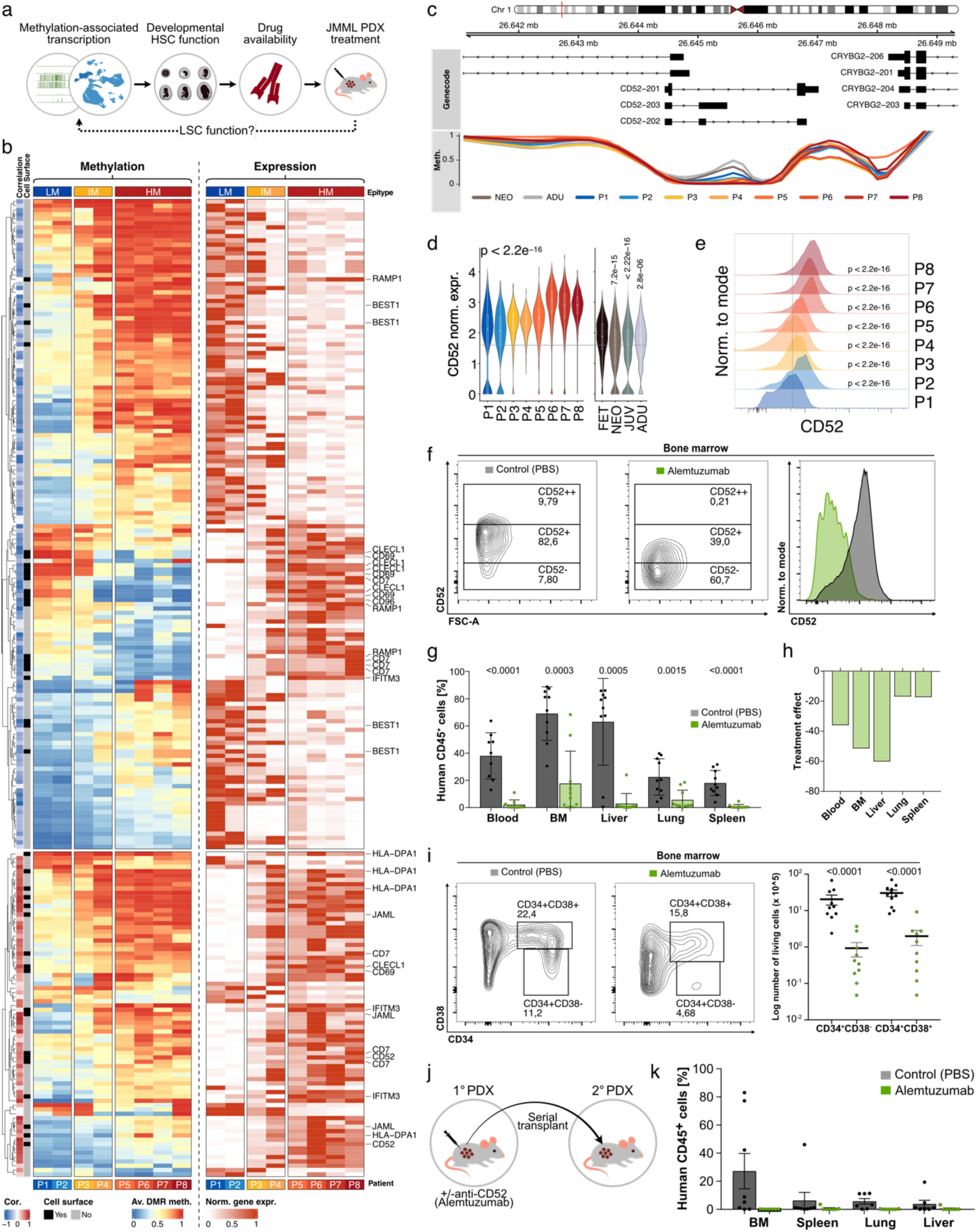
Integrative analysis identifies novel prognostic biomarkers and therapeutic targets for HM-JMML. **a**, Strategy for the identification of disease-specific candidate genes in JMML HSCs. **b**, Identification of methylation-associated gene expression changes across JMML epitypes. The heatmaps display expression (right) and methylation (left) values of the DMR-DEG associations for the 56 high-confidence candidate genes identified using a generalized linear model (for details, see Material and Methods). Normalized gene expression was scaled to the minimum and maximum for each gene. For each DMR the average beta values across all CpGs per region are depicted. ‘Correlation’ depicts the Pearson correlation coefficients as per DMR-DEG association. ‘Cell surface’ indicates if a gene is predicted to be expressed on the cell surface. The 13 genes encoding for cell surface proteins are indicated by name. Of note, a single gene can be associated with multiple DMRs. **c**, Locus line plot showing the smoothened methylation beta values of the *CD52* locus in HSCs across JMML patients and healthy references. **d**, Violin plots showing the normalized expression profiles of *CD52* in JMML and healthy reference HSCs. Significance of expression differences across samples was tested using Kruskal-Wallis test. Significance of the expression differences of postnatal (i.e. NEO, JUV, ADU) to prenatal (FET) HSCs was tested using Wilcoxon rank sum test. Boxplot center line corresponds to data median, box height to the inter-quartile range and lines to lower / upper quartile +/− 1.5 times inter-quartile range. **e**, Histograms showing the cell surface expression of CD52 in Lin^-^CD34^+^CD38^-/lo^ JMML HSPCs using flow cytometry analysis of patients across epitypes. Significance was calculated per patient with Wilcoxon test against P1. **f-k**, Patient-derived xenograft (PDX) mouse experiments. For experimental details refer to ‘Materials and Methods’ and Extended Data Fig. 12a. **f**, Representative FACS contour plot and histogram showing CD52 expression on engrafted human CD45^+^ JMML cells in mice treated with alemtuzumab (green) or with vehicle control (dark grey). Expression levels are depicted on a logarithmic scale. **g**, Quantification of engrafting human CD45^+^ cells after 4 weeks of treatment with alemtuzumab (green) or vehicle (dark grey). Depicted are the human cells as a percentage of all hematopoietic cells detected in each organ tested. **h**, Effect of alemtuzumab treatment (‘treatment effect’) depicted as the differences of the mean engraftment in treated vs untreated mice per organ (treatment effect = mean_treated_ – mean_untreated_). **i**, *Left:* representative FACS plot showing CD34 and CD38 expression on human CD45^+^Lin^-^ JMML cells engrafted in the bone marrow of mice treated with alemtuzumab (green) or with vehicle control (dark grey). *Right:* Quantification of Lin^-^CD34^+^CD38^-^ and Lin^-^CD34^+^CD38^+^ human HSPCs. Depicted is the logarithm of the absolute number of living human cells per femur. **j**, Schematic depicting the 2° transplantation experiments. Secondary recipient mice received bone marrow cells from alemtuzumab-treated or vehicle-treated primary PDX mice. **k**, Frequency of human CD45^+^ cells in 2° recipients of cells from alemtuzumab-treated (green) or vehicle-treated (dark grey) donor mice. Engraftment was evaluated 6 weeks after transplantation.

Focusing on HM-JMML as the epitype with the most urgent need for novel therapeutic approaches, we identified *CD52*, which encodes a cell surface protein that is an established drug target in chronic lymphocytic leukemia and multiple sclerosis^29–31^. *CD52* is associated with epigenetic aberrations in JMML HSCs (Fig. 6b,c), and shows moderate expression levels in healthy HSCs across normal hematopoietic maturation, with a slight but significant upregulation in fetal HSCs (Fig. 6d). However, it is strongly upregulated in JMML HSCs with the highest expression levels found in HM-JMML (Fig. 6d). Epitype-specific CD52 expression was confirmed at the protein level in CD34^+^CD38^-/lo^ HSPCs from JMML patients, suggesting CD52 surface protein as an accessible drug target on JMML HSCs (Fig. 6e). Finally, murine *Cd52* was transcriptionally upregulated in HSCs of *Ptpn11^E76K/+^* mice relative to control HSCs, confirming the disease-specific upregulation in JMML HSCs (Extended Data Fig. 11d). In conclusion, CD52 is aberrantly upregulated preferentially in HM-JMML and is a promising target for further evaluation as a prognostic biomarker and as a therapeutic target in HM-JMML.

### Anti-CD52 treatment reduces leukemia burden, depletes leukemic stem cells, and disrupts disease propagation in a JMML PDX model

To assess the therapeutic potential of anti-CD52 targeted treatment, we used an established preclinical patient-derived xenograft (PDX) mouse model of JMML and treated PDX mice with the therapeutic monoclonal antibody alemtuzumab for 4 weeks (Extended Data Fig. 12a, Supplementary Table 20)^32,33^. FACS analysis of primary recipients revealed an efficient depletion of human CD52^+^ cells following alemtuzumab treatment, resulting in a significantly reduced frequency of human leukemia cells in blood, bone marrow, spleen, liver, and lung (Fig. 6f-h and Extended Data Fig. 12b,c). As proposed by our integrative multi-omics analysis, alemtuzumab treatment targeted human HSPCs as evidenced by an efficient depletion of Lin^-^ CD34^+^CD38^-^ cells (Fig. 6i). This resulted in the depletion of virtually all human hematopoietic cells in our xenograft model, including CD52^-^ cells (Extended Data Fig. 12c-h). Secondary transplantation of total bone marrow from alemtuzumab- or vehicle-treated animals showed that anti-CD52 treatment prevented leukemic engraftment in 2° recipients (Fig. 6j,k). In conclusion, therapeutic targeting of CD52 leads to efficient depletion of leukemia-propagating stem cells *in vivo*.

## Discussion

In the present study, we performed a multi-layered molecular analysis in which we systematically compared JMML across disease-specific epitypes to healthy references of HSC development. Single-cell transcriptome and DNA methylome analyses gave opposing results: transcriptomes revealed pronounced fetal HSC signatures for the more aggressive JMML epitypes whereas DNA methylomes indicated a postnatal epigenomic pattern in all patients analyzed. This finding is particularly important as the current consensus assumes that early childhood malignancies arise during fetal development and manifest as maturation blocks in aberrantly developing tissues^2,11,34^. This view is supported by a number of studies which provide mostly genetic evidence for fetal origins, which are based on the detection of driver mutations in neonatal blood spots or on mathematical modeling of the originating developmental stage based on cell cycle-related mutation rates^2,21,35–39^. However, whether or not the presence of fetal-like transcriptional programs present across a range of childhood malignancies can unambiguously be considered as indicative of their prenatal origin remains a matter of scientific debate^4,40–43^. In the present study, an in-depth analysis of the transcriptome layer revealed that malignant hematopoietic stem cells (HSC) hijack transcription programs across different developmental stages and that HSCs from high-risk patients reactivate fetal-like gene expression signatures in a patient-specific manner. To complement the transcriptomic layer with epigenetic information, we performed DNA methylome analysis, which presumably provides more robust information on developmental and tissue origins of tumors than transcriptomes, as multiple lines of evidence suggest that DNA methylomes represent the developmental and differentiation states in which the malignant cells have been arrested^26,44–48^. In the context of JMML, DNA methylome analysis unambiguously showed postnatal epigenomes in HSCs of all JMML samples investigated in this study by phylogeny and epigenetic scar analyses. This indicated that fetal-like transcription programs observed in JMML are unlikely to be caused by a developmental maturation block.

These apparently conflicting results might be explained by OFR, a phenomenon well-known in the context of adult but not childhood malignancies. OFR describes the re-acquisition of fetal gene expression programs in tumorigenesis, which has been described for adult malignancies such as hepatocellular carcinoma^1^. In general, the process of oncogenesis requires cellular reprogramming to sustain cells with physiologically contradicting features such as stemness/pluripotency and differentiation bias^49^. Thus, OFR might reflect a tumor-specific strategy to gain cellular plasticity and to overcome physiological limitations of normal development. In JMML, the OFR hypothesis is supported by the clinical observation that high-risk JMML patients are typically older than 2 years at the time of diagnosis^13,16^. In addition, a recent publication tested JMML driver mutations in Guthrie cards from individuals later diagnosed with JMML and found that 9/9 patients with low-risk (i.e. LM epitype) JMML had a detectable RAS-pathway mutation at birth, while only 3/7 patients with higher risk (i.e. IM or HM epitype) JMML were tested positive for a JMML driver mutation at birth^21^. Of note, whereas the current literature suggests that LM JMML arises prenatally, our data reveals postnatal maturation signatures at both the transcriptomic and epigenomic levels. This underlines that LM JMML, although most likely arising prenatally, does not show fetal signatures indicative of a maturation block. Vice versa, all HM patients analyzed here revealed postnatal epigenomes combined with pronounced enrichment of fetal-like transcription signatures, supporting the OFR hypothesis. It remains to be studied how many patients ultimately acquire their disease-initiating mutation postnatally and are affected by OFR. Furthermore, it is unclear whether OFR is influenced by the type of driver mutation.

Since multi-layer molecular analyses from primary patient samples are descriptive and cannot prove causality, we used a JMML mouse model to test whether the expression of a JMML driver mutation in young adult HSCs is sufficient to establish a fetal-like HSC gene expression program. Indeed, we found that induction of a *Ptpn11^E76K^* mutation in postnatal HSCs was sufficient to establish an aberrant fetal-like transcription program, hence formally proving that OFR can occur in JMML. In summary, our work highlights how a multi-layered molecular analysis together with experimental validation enable the dissection of the characteristics of distinct disease phenotypes within a uniformly diagnosed and genetically characterized disease entity such as JMML. Our work, demonstrates that OFR is a phenomenon with relevance to childhood malignancies and might motivate the re-evaluation of the assumed pathogenetic mechanisms in relation to embryonic or fetal development across the entire spectrum of childhood malignancies. Nevertheless, there are still open questions that remain to be answered in the future: 1) How are the transcriptional and epigenetic aberrations established? 2) What is the order of acquisition of aberrations, i.e. is the transcriptome or the epigenome altered first? 3) How can the observed phenotypic heterogeneity be explained in the absence of genetic heterogeneity (i.e. presence of the same disease-initiating mutation)? Although answering these questions is beyond the scope of the current manuscript, it is tempting to speculate that transcriptional re-programming might precede the epigenetic alterations seen in the context of JMML. This hypothesis would be supported by the observation that, despite the presence of fetal-like transcription programs, the DNA methylomes lack pronounced signs of fetal reprogramming in the JMML mouse model (data not shown). Along these lines, DNA hypermethylation has been associated with increased proliferation activity and in some cases with methylation age^50–53^.

The discovery of OFR highlighted that JMML HSCs express a complex mosaic of expression programs across developmental stages. As a consequence, it is important to consider healthy references across developmental stages to identify cancer-specific aberrations. This approach provided the basis for the identification of prognostic biomarkers and novel therapeutic targets for this subset of patients with particularly poor outcome. For example, we could demonstrate that aberrant DNA methylation is accompanied by elevated mRNA expression levels of *CD52* in leukemic HSCs, especially in HSCs of high-risk JMML patients. This elevated *CD52* mRNA expression was not only recapitulated in HSCs of our JMML mouse model but it also translated into elevated surface protein expression levels on patient HSCs. This raises the possibility of performing clinical risk-stratification of JMML patients using a flow cytometry-based assay without the need for specialized and time-consuming epigenetic analyses. Of note, *CD52* has recently been described to play a functional role in both pre- and postnatal HSC development^27,49,54,55^. Moreover, in line with our observation of an early differentiation bias in JMML HSCs, CD52 has been implicated in early myeloid priming^56,57^. A therapeutic monoclonal antibody targeting CD52 (alemtuzumab) is approved by the FDA and EMA for the treatment of multiple sclerosis^29,58,59^ and has been used as part of conditioning regimens for allogeneic hematopoietic stem cell transplantation (HSCT)^60,61^. This made CD52 an interesting candidate for pre-clinical target validation, as HSCT is the only potentially curative treatment for high-risk JMML, although, more than 50% of patients relapse post-HSCT^62^. Furthermore, pre-clinical therapeutic activity of an approved drug could potentially be rapidly translated into the clinical setting. Using a patient-derived xenotransplantation model of JMML, we observed an efficient depletion of CD34^+^CD38^-^ JMML hematopoietic stem and progenitor cells upon alemtuzumab treatment, which was functionally evidenced by the depletion of virtually all human hematopoietic cells independent of cell type-specific CD52 expression levels. Furthermore, alemtuzumab treatment abrogated the engraftment of human JMML cells in secondary recipient animals. Of note, this therapeutic effect was observed in PDX mice of the high-risk patient P7, who clinically showed a relapse after allogeneic HSCT. Hence, CD52-targeted therapy with alemtuzumab shows promising pre-clinical efficacy in high-risk JMML. This provides a rationale to further assess anti-CD52 treatment in pre-clinical studies as an option for JMML patients with high risk of relapse after allogeneic hematopoietic stem cell transplantation, for example in the context of the conditioning regimen.

In conclusion, our work on JMML provides evidence for oncofetal reprogramming as a mechanism for the expression of fetal-like transcriptional programs in a childhood malignancy. Furthermore, we demonstrate how a molecular high-precision approach identifies aberrantly regulated developmental markers and guides the identification of disease biomarkers and novel targeted therapies for a notoriously difficult-to-treat childhood leukemia.

## Methods

### Authorization and ethical approval

The study was conducted according to the Declaration of Helsinki after written informed consent was obtained from the healthy subjects, the patients or their legal guardians. All experiments were approved by the local institutional review boards. All information is subject to the provisions of the local data protection regulations. JMML samples were obtained from the Hilda Biobank Freiburg, and fetal samples were obtained from the Wellcome Trust-funded Human Developmental Biology Resource (HDBR; http://www.hdbr.org). HDBR is regulated by the UK Human Tissue Authority (HTA; www.hta.gov.uk) and operates in accordance with the relevant HTA Codes of Practice. Healthy BM donors at the Heidelberg University Hospital received financial compensation. All mouse experiments were approved by German local authorities (Regierungspräsidium Karlsruhe and Regierungspräsidium Freiburg). All data were stored in accordance with the DKFZ framework for data protection.

### Isolation of hematopoietic cells from spleen of JMML patients

Splenectomy specimens from patients diagnosed with JMML were used to prepare single cell suspensions followed red blood cell lysis. To obtain mononuclear cell (MNC) fraction, single cell suspensions prepared from spleen tissue were subjected to density gradient centrifugation (Ficoll). Mononuclear cells were viably frozen and used for downstream analysis.

### Primary human pediatric bone marrow

Healthy juvenile pediatric bone marrow samples for WGBS were obtained from children at the age of 0 to 7 years in the course of neurosurgical interventions (e.g., correction of craniosynostoses, craniotomies performed for epilepsy surgery) after written informed consent was obtained from the patients’ legal guardians. For this purpose, the diploe of the craniotomized bone was cannulated with 27G needles and manually flushed with Ringer’s solution. Downstream procedures for sample preservation were identical as for materials obtained from JMML patients. For scRNA-seq, left-over bone marrow mononuclear cells from pediatric healthy normal sibling donors were isolated by ficoll gradient centrifugation.

### Primary human fetal spleen and liver

Tissue from healthy prenatal references was obtained from the HDBR. Developmental age, sex, and aneuploidy screens were performed as previously described^63^. Liver (n = 4; 12-16 post conception weeks (PCW)) and spleen (n = 3; 12-16 PCW) samples were diced and digested with 1.6mg/mL collagenase type IV (Worthington) in RPMI (Sigma-Aldrich) supplemented with 10% (v/v) heat-inactivated fetal bovine serum (Gibco), 100 U/mL penicillin (Sigma-Aldrich), 0.1 mg/mL streptomycin (Sigma-Aldrich), and 2 mM L-Glutamine (Sigma-Aldrich) for 30 min at 37°C with intermittent shaking. Digested tissue was passed through a 100 μm filter, and cells collected by centrifugation (500 g for 5 min at 4°C). Cells were treated with 1X RBC lysis buffer (eBioscience) for 5 min at room temperature and washed once with flow buffer (PBS containing 5% (v/v) FBS and 2 mM EDTA) prior to counting. Cells were cryopreserved in 90% (v/v) FBS with 10% dimethylsulfoxide (DMSO) to permit analysis in batches.

### Primary human bone marrow from adults

As healthy adult references, human samples from healthy adult individuals at the age of 23 to 25 were collected from bone marrow aspirates of healthy donors via iliac crest. Mononuclear cells were isolated by Ficoll (GE Healthcare) density gradient centrifugation and stored in liquid nitrogen until further use.

### Single-cell RNA-seq and WGBS of JMML patients

For scRNA-seq and ultra-low input whole-genome bisulfite sequencing (WGBS) of human JMML spleen and pediatric BM samples, cells were thawed in a water bath at 37°C and transferred dropwise into prewarmed IMDM (Gibco) supplemented with 10% FCS and 2 U/ml DNase I. Cells were centrifuged for 5 min at 300 g and washed with PBS (Gibco) supplemented with 4% FCS and DNase I (LIFE Technologies). Cells were resuspended in PBS containing 4% FCS (4 million per 100 µl) and fluorophore-conjugated antibodies for lineage markers, CD34, CD38, CD45RA, and CD90 (Supplementary Table 21), followed by incubation for 30 min at 4 °C in the dark. Cells were washed with PBS containing 4% FCS and resuspended in an appropriate volume for flow cytometry. For live/dead staining, 4,6-diamidino-2-phenylindole (DAPI; 1:1,000) was supplemented to the cell suspension. After 3 min of incubation at room temperature, cells were filtered through a 40 μm cell strainer (Greiner). Cell sorting was performed using BD FACSAria Fusion. For scRNA-seq of JMML HSPCs, 1,700 to 10,000 Lin^-^CD34^+^CD38^dim/-^ cells were sorted into IMDM (Gibco) containing 4% FCS and directly subjected to droplet-based scRNA-seq (3’ v2 protocol from 10X Genomics). All scRNA-seq libraries from primary JMML patient material were sequenced on a HiSeq4000 (Illumina) paired-end 26+74 bp at the DKFZ Genomics & Proteomics Core Facility (GPCF) (Heidelberg, Germany). For WGBS of JMML Lin^-^CD34^+^CD38^dim/-^ HSPCs, 33 to 250 cells were directly sorted into RLT plus lysis buffer and stored at −80°C for later use.

### Single-cell RNA-seq and WGBS of healthy pediatric references

For WGBS of healthy juvenile reference bone marrow, cells were isolated from frozen samples as described above for JMML and 500 Lin^-^CD34^+^CD38^-/lo^CD45RA^-^CD90^+^ HSCs were directly FACS-sorted into RLT plus lysis buffer. For WGBS of prenatal HSCs from fetal liver and spleen, up to 1 million cells were stained with antibody cocktail (Supplementary Table 22), incubated for 30 min on ice and washed with flow buffer. Cells were resuspended at 10 million cells per ml and DAPI added immediately before FACS (Sigma-Aldrich; final concentration of 3 μM). Live, single, Lin-, CD34^+^CD38^-/lo^CD45RA^-^ CD90^+^ cells were sorted into RLT plus lysis buffer. Cells were stored at −80°C for later use. For scRNA-seq of healthy pediatric bone marrow, mononuclear cells were enriched for immature compartments using CD34^+^ progenitors and Lin^-^CD34^+^CD38^-^ HSPCs using flow cytometry. Viable live gate, progenitor- and stem cell-enriched populations were subjected to chromium 10X processing using v3 chemistry (10X Genomics).

### WGBS of healthy adult references

For WGBS of adult stem and progenitor cells, cryopreserved adult bone marrow samples were thawed and stained in FACS buffer (FB) (PBS supplemented with 5% FCS and 0.5 mM EDTA) containing the respective antibodies (Supplementary Table 23) for 30 minutes followed by washing with FB. Subsequently, cells were sorted using an BD Aria Fusion II sorter. HSPC populations were sorted as previously defined^64^: Lin^-^CD34^+^CD38^-/lo^CD45RA^-^CD90^+^ HSCs, Lin-CD34+CD38-CD45RA-CD90-MPPs, Lin-CD34+CD38+CD10-CD45RA-CD135+ CMPs, Lin-CD34+CD38+CD10-CD45RA+CD135+ GMPs, and Lin-CD34+CD38+CD10-CD45RA-CD135-MEPs. WGBS libraries were generated using a published tagmentation-based protocol^65^.

### Flow cytometry analysis of JMML patients

For validation experiments of epitype-specific CD45RA/CD90 expression profiles, single cell suspensions prepared from JMML spleen tissue were subjected to density gradient centrifugation (Ficoll) to obtain mononuclear cell fractions (MNCs). An aliquot of the isolated MNCs was stained for flow cytometry (Supplementary Table 24) and data acquired using BD LSRFortessa.

For validation experiments of epitype-specific CD52 expression profiles, JMML samples were thawed as described above (“Single-cell RNA-seq and WGBS of JMML patients”) and stained with fluorescence-coupled antibodies for flow cytometry (Supplementary Table 25). Cells were analyzed with BD FACSAria Fusion.

### Panel-seq for JMML genotyping

DNA from bone marrow granulocytes was used for amplicon-based, next-generation deep sequencing to determine the variant allele frequencies (VAFs) of leukemia-specific index mutations. Targets of JMML associated Genes were enriched using a custom in-house panel (AmpliSeq JMML panel v2, IAA11909_192; Thermo Fisher Scientific) and processed with the NEBNext Ultra II DNA library preparation kit (New England Biolabs). Sequencing was performed on MiSeq sequencers (Illumia). Sequence variants were evaluated according to the ACMG classification system.

### Processing of scRNA-seq data

scRNA-seq data from JMML patients and JUV references were generated in this study, data from FET, NEO, and ADU references were obtained from the Human Cell Atlas project (Supplementary Table 2). Sequencing data from JMML patient samples for scRNA-seq were aligned against the hg38 human reference genome, and gene-cell matrices were generated using *CellRanger* (v.2.1.1). Subsequent data processing included the sample-wise inclusion of cells that met specific quality criteria:

1.) The number of genes exceeded 100.
2.) The number of genes was within three median absolute deviations (MADs) from the median, determined using the “isOutlier” function in *scater* (v.1.10.1).
3.) The number of counts exceeded 200.
4.) The number of counts was within three MADs from the median, determined using the “isOutlier” function in *scater* (v.1.10.1).
5.) Mitochondrial content was required to be below a 5% cutoff.

The ADU dataset was annotated to 35 cell types according to Hay et al., 2018^66^. Subsequently, the annotated ADU dataset was down-sampled to a maximum of 2500 cells per cell type and served as a label transfer reference to annotate other datasets, including FET, NEU, JUV, and JMML. Label transfer was performed using the “FindTransferAnchors” and “TransferData” functions (dims = 1:30) in *Seurat* (v.3.1.5)^67^. Each cell was assigned one transferring score for each cell type, and the cell type assignment of cells was based on the highest cell type transferring score.

In the FET dataset, cells with annotations of endothelial cells, fibroblasts, hepatocytes, Kupffer cells, and VCAM1+ EI macrophages, based on the original paper^68^, retained their original annotations.

For the healthy reference datasets, each cell type was down-sampled to a maximum of 2500 cells for further analysis.

### scRNA-seq data integration

We employed the “anchor” integration method to integrate scRNA-seq datasets from JMML patients and healthy references using *Seurat* (v.3.1.5). The following steps were taken with default settings:

1.) Dataset-wise data normalization was performed using the “NormalizeData” function.
2.) Dataset-wise feature selection was conducted with the “FindVariableFeatures” function.
3.) Pairwise “anchors” between the datasets were identified using the “FindIntegrationAnchors” function (dims = 1:30).
4.) Data integration based on these “anchors” was achieved through the “IntegrateData” function (dims = 1:30).
5.) The integrated data underwent regression of counts, scaling, and centering using the “ScaleData” function, followed by PCA dimensionality reduction.

The scRNAseq data of CD34+-enriched HSPCs and MNCs from JMML patients were integrated using the ‘anchor’ integration methods in the Seurat (v.3.1.5). Similar to the integration of JMML with healthy references, patient-wise data normalization and feature selection were performed. Pairwise ‘anchors’ between patients were identified and integrated. The integrated data was scaled and PCA calculated.

### Pseudotemporal analysis of hematopoietic lineages

PCA was used to estimate a linear latent pseudotime for each cell across distinct hematopoietic lineages using RNA expression data.

RNA expression data was normalized by dividing the feature counts of a given cell by the total counts for that cell, then multiplying the quotient by 10.000 and finally log1p transforming the product (default settings for *Seurat* “NormalizeData”).

To facilitate PCA in capturing lineage-specific changes, we devised a gene selection procedure consisting of three main steps:

1.) The dataset was restricted to non-tumor cells, and genes with no observed transcripts across all cells were excluded.
2.) For each gene, a cumulative link model was constructed to establish a relationship between the position of cells within the lineage and their normalized expression of the respective gene.
3.) Genes satisfying both a Benjamini-Hochberg adjusted model coefficient p-value less than 0.01 and an absolute coefficient value greater than 1.5 were selected.

In the case of the dendritic cell lineage, this procedure was executed using 10,949 non-tumor cells, leading to the identification of 2,205 genes. Subsequently, the normalized expression values of each gene were scaled by subtracting the per-cell expression values by the average expression of the gene and dividing the difference by its standard deviation. This scaling process adhered to the default settings for the *Seurat* “ScaleData” function. The scaled expression values were then employed to perform PCA using *Seurat*.

Spearman correlation was calculated to evaluate the concurrence of differences in PCA cell embeddings and various metadata features. For categorical metadata, one-hot encoding was utilized to compute correlations. If necessary, for visualization purposes, the principal component with the highest correlation with lineage was flipped so that the lineage commenced on the left-hand side of the plot.

### UMAP integration of scRNA-seq data from CD34^+^-enriched JMML HSPCs and JMML MNCs

To project the scRNA-seq data of FACS enriched JMML CD34^+^CD38^-/lo^ HSPCs onto the existing MNC UMAP space we used the MNC scRNA-seq set as a reference and identified anchors between both datasets by applying the *Seurat* “FindTransferAnchor” function with PCA as a specified reference reduction. We mapped the HSPC query data onto the MNC UMAP space using the “MapQuery” function and visualized the combined UMAP using *ggplot2*^69^.

### Transcriptional priming and leukemia-related transcription programs of human HSCs

Gene expression signatures were extracted from previous publications ^57,70,71^. Transcriptional priming was calculated using the *Seurat* “AddModuleScore” function for 1000 randomly selected HSCs from each epitype. Statistical significance was determined based on two LMMs as implemented in the *lme4* R package followed by ANOVA:

LMM1 = signature ∼ Subgroup + (1|Donor)

LMM2 = signature ∼ (1|Donor)

### Differential gene expression analysis

Differentially expressed genes (DEGs) between JMML HM and JMML nonHM CD34+enriched samples were identified using the “FindMarkers” function with default settings in *Seurat* (v.3.1.5). A total of 255 DEGs with an adjusted p-value < 0.05 and an absolute log fold-change < 0.25 were obtained.

Developmental stage-specific DEGs were identified in the cells annotated as HSCs in four healthy reference datasets (FET, NEO, JUV, and ADU) using the “FindAllMarkers” function in *Seurat* (v.3.1.5) for one-vs-rest comparisons. In each stage, the top 50 upregulated DEGs with an adjusted p-value < 0.05 and an absolute log fold-change < 0.25 were obtained. Any overlapping DEGs across stages were removed. We identified 44 DEGs for FET HSCs, 41 DEGs for NEO HSCs, 50 DEGs for JUV HSCs, and 47 DEGs for ADU HSCs.

### Enrichment of transcription signatures using Metascape

For functional gene list analyses, *Metascape*^72^ was used applying default settings. As input, up- or downregulated HM vs nonHM DEGs of JMML HSCs were used (Supplementary Table 7).

### Logistic regression analysis of scRNA-seq data

The logistic regression method was used to measure the transcriptional similarity between JMML HSCs and healthy reference cell types (FET, NEO, JUV, and ADU) across developmental stages, as previously described ^5^. In brief, we first trained a logistic regression model on the scRNA-seq count matrix of healthy references, encompassing cells of all hematopoietic cell types, using the trainModel function with default settings in LogisticRegression.Core.R. This trained model was then applied to predict the likelihood of similarity between cell types in healthy references and JMML patients using the predictSimilarity function with minGeneMatch = 0.5. The resulting average similarity scores of HSCs in JMML patients against all cell types were visualized in a heatmap using the ComplexHeatmap^73^ package.

### DNA methylation array analysis

For DNA methylation array profiling, gDNA of 200,000 MNCs per sample was extracted and submitted to the DKFZ GPCF (Heidelberg, Germany) for Infinium MethylationEPIC BeadChip (Illumina) analysis. Data processing and epitype prediction were performed as described previously^16^.

### Library preparation for ultra-low input whole-genome bisulfite sequencing (WGBS)

For ultra-low input WGBS library preparation, we used max. 500 cells and followed a customized previously published single-cell Bisulfite Sequencing (scBS-seq) protocol ^74,75^. In brief, we sorted cells using flow cytometry into RLT Plus buffer to a total volume of 10 μL, which was applied to the following changes: single preamplification for 90 minutes at 37°C; 14 cycles of library amplification; two time 0.7x SPRI selection for library purification. Molarities and fragment sizes were measured and libraries were submitted for Illumina sequencing on a HiSeq X with paired-end 150 bp and 10% PhiX to the DKFZ GPCF.

### Processing of whole-genome bisulfite sequencing data

The Omics IT and Data Management Core Facility (DKFZ, Heidelberg) processed WGBS data as described earlier^76^. In short, reads were aligned using an updated pipeline published by Wang et al.^65^, implemented as a Roddy Workflow (https://github.com/DKFZ-ODCF/AlignmentAndQCWorkflows) in the automated One Touch Pipeline^77^. In short, adaptor sequences of raw reads were trimmed using *Trimmomatic*^78^. Next, sequencing reads were in silico bisulfite-converted^79^(C>T for the first read in the pair, G>A for the second). *BWA-MEM*^79^ was used with default parameters to align the converted reads to the in silico bisulfite-converted reference genome hg19, extended with the PhiX and lambda phage sequences. Post alignment, reads were converted back to their original state. Reads with a mapping quality ≥25 and nucleotides with a Phred-scaled quality score ≥25 were considered for PCR duplicate removal per sequencing library using the software *Picar*^80^. Methylation calling and M-bias QC was performed using *MethylDackel* ^81^ v0.4.0 and the parameters *--OT 6,146,2,144 --OB 7,146,12,150*, according to the M-bias quality control plots. Methylation calls were imported into R^82^ v3.6 using the R package *methrix*^83^ v 1.0.05, summarized based on annotated reference indices, and collapsed based on strand information. Furthermore, single nucleotide polymorphisms (SNP) with a minor allele frequency >0.1 were filtered, using the reference provided in the R package *BSgenome.Hsapiens.UCSC.hg19*^84^ v 1.4.0, and initial quality control was performed. For downstream analysis, DNA methylation data were imported into the R package *bsseq*^85^ v1.20.0 and technical replicates were collapsed. To define average CpG island (CpGi) methylation, CpGi annotation was extracted using the R package *RnBeads*^86^ v 2.4.0. For principal component analysis (PCA), methylation levels of the 20,000 most variable methylated CpGs were applied to the R base function *prcomp*.

### Methylation-based cell type classifier

For DNA methylation-based cell type classification, a previously published algorithm was applied^24^. As normal references, WGBS data of adult Lin^-^CD34^+^CD38^-/lo^CD45RA^-^CD90^+^ HSCs, Lin^-^CD34^+^CD38^-/lo^CD45RA^-^CD90^-^ MPPs, Lin^-^CD34^+^CD38^+^CD10^-^CD45RA^-^CD135^+^ CMPs, Lin^-^CD34^+^CD38^+^CD10^-^CD45RA^+^CD135^+^ GMPs, and Lin^-^CD34^+^CD38^+^CD10^-^CD45RA^-^ CD135^-^ MEPs was employed, which was generated in course of this study.

As input to the classifier, we used WGBS data that was prepared as outlined in the previous section. Briefly, WGBS data was subsetted to regions outlined in Farlik *et al*.^24^, using the hg38 to hg19 liftover function from the R package *rtracklayer*^87^. The average methylation within each region was computed and used to execute the “cellTypePredictor.R” script by Farlik *et al*.^24^. The classifier itself was trained on normal references and applied to JMML tumor samples.

### Differential methylation analysis

To define differentially methylated loci, a linear model was applied, considering the epigenotype (HM and non-HM) and sorting quadrant of the respective sample as a covariate. Therefore, the function “DMLfit.multiFactorR” with the parameter, *smoothing = TRUE*, from the R package *DSS*^88^ v2.32.0 was applied. The epigenotype HM coefficient was extracted using the function “DMLtest.multiFactor” and differentially methylated regions (DMRs) were defined using the function “callDMR”, based on regions with >3 CpGs, a length of >50 bps, and a Benjamin-Hochberg corrected *P* value <0.05 in at least 50% of all CpGs within that region. Additionally, DMRs were subsetted for an average coverage of ≥5 in at least half of the samples belonging to the HM and non-HM epigenotype respectively and a difference in methylation >0.2. DMRs were annotated using the R package *ChIPseeker*^89^ v1.31.3.900 and *TxDb.Hsapiens.UCSC.hg19.knownGene*^90^ v3.2.2. Hierarchical cluster analysis of DMRs was performed using the R package *pheatmap*^91^ v1.0.12, applying “ward.D2” as the clustering method to “Manhattan” distances.

### Transcription factor motif enrichment

DNA transcription factor (TF) motif enrichment was performed using the command line tool *Homer*^92^ (v4.9.1) and the parameter *-size given*. DMRs were stratified in hypo- and hypermethylated regions, and a set of random regions with equal size, similar CpG frequency (15% tolerance in the deviation of CpG composition were allowed), and a maximum of 20% N’s in the sequence composition, were used as a background. The top 10 most significantly enriched TF motifs in hypo- and hypermethylated DMRs were visualized.

### Locus overlap and enrichment analysis of ChromHMM states

The R package *LOLA*^93^ (v1.16.0) was used to enrich DMRs with ChromHMM states acquired from Encode^94^. DMRs were stratified in hypo- and hypermethylated regions and enriched to a background containing a set of random regions with equal size, similar CpG frequency (15% tolerance in the deviation of CpG composition was allowed), and a maximum of 20% N’s in the sequence composition.

### Locus plots

*IGV* (v2.12.2) was used to visualize locus-specific methylation at single CpG resolution of WGBS and DNA methylation array data (450k). Alternatively, the R package *gviz*^95^ v1.30.3 was used to generate locus plots. For the visualization of WGBS data, methylation calls were smoothed with the R package *bsseq*^85^ v1.20.0, applying the function “BSmooth” with default parameters.

### Homeobox gene classes

Homeobox genes were taken from Holland *et al.* ^96^. We annotated DMRs obtained from comparing HM vs non-HM JMML patients using the “annotatePeak” function from the *ChIPseeker* package^89^, assuming a tssRegion of +/− 3,000 base pairs. Subsequently, we counted how many DMR associated genes were present in either of the previously defined Homeobox gene classes PRD, ANTP and Misc (LIM, POU, HNF, SINCE, TALE, CUT, PROS, ZF, CERS). We divided the size each gene set by 30,000, and the number of DMR associated genes occurring in each gene set by the number of total DMR associated genes. The log2 fold change of those ratios was plotted.

### Delineating methylation profiles linked to healthy hematopoietic stem cell development and maturation

To analyze JMML within the context of HSC development, we compared the methylomes of JMML HSCs to those of healthy fetal, neonatal, juvenile, and adult HSCs. For this purpose, the entire genome was segmented into contiguous 1kb regions, and the average methylation in regions covered in all samples was computed (n = 1,209,679). Average methylation within each region was subsequently correlated with the developmental order, progressing from fetal to neonatal, juvenile, and adult stages, using spearman correlation.

Of all the regions analyzed, 1,608 exhibited an absolute Spearman correlation coefficient of at least 0.9, corresponding approximately to the 0.999 quantile of the absolute Spearman correlation distribution.

As the methylomes of fetal and juvenile samples were measured using post-bisulfite adaptor tagging (PBAT) while those of neonatal and adult samples were assessed by TWGBS, there remained a residual risk of confounding by the sequencing method. To address this issue, we computed the Spearman correlation of average methylation and sequencing method, subsequently removing all regions with an absolute correlation coefficient greater than 0.4 to account for the sequencing method covariate. This threshold corresponded approximately to the 0.75 quantile of the absolute Spearman correlation distribution. The remaining 1,179 regions were presumed to capture the epigenetic changes occurring throughout HSC development

For the reconstruction of phylogenetic trees representing HSC development, we calculated the Manhattan distance between samples based on the average methylation of the selected 1,179 regions. Tree reconstruction, assuming minimum evolution, was conducted using the “fastme.bal” function from the R package *ape*^97^ (version 5.6.2).

### Principal component analysis of JMML and healthy reference bulk methylomes

To focus on the HSC bulk methylomes, data were subset accordingly, with the exclusion of the FL2_HSC_PBAT_2 sample due to its poor quality. The methylation values of the 10,000 most variable CpGs across all remaining samples were centered around 0 and scaled by their standard deviation. Subsequently, Principal Component Analysis (PCA) was applied to the scaled methylation values.

PCA embeddings and variance explained per principal component were computed using the prcomp function in base R, thereby facilitating an unsupervised analysis of genome-wide methylation patterns across hematopoietic stem cells.

### Identifying epigenetic scars of development

Epigenetic scars, conceptually, are characterized by marked methylation changes between distinct developmental states, while remaining stable during subsequent development (related to Abdel-Hakeem et al.^98^). To identify epigenetic scars, DMRs were called between a developmental state of interest and all succeeding states. “callDMR” from the R package *DSS*^99^ (2.42.0) was used to identify DMRs across two groups. The “callDMR” function was invoked using the following arguments: delta=.1, p.threshold=.05, minlen=50, minCG=3, dis.merge=50, pct.sig=0.5. Identified DMRs were filtered to suffice an average coverage of five in at least 50% of samples in both groups.

We identified 3,332 potential scars from comparing prenatal to postnatal states, and, 8,964 from the comparison between neonatal and post-neonatal states. Given the lenient nature of the DMR calling procedure, only a small fraction of all DMRs represent genuine epigenetic scars.

To select DMRs consistent with the concept of epigenetic scars, we applied the following filtering approach:

1.) Conduct a Fisher’s exact test on a 2×2 contingency table comparing the total number of methylated and unmethylated CpGs within a given DMR and the two groups being compared. DMRs meeting a Benjamini-Hochberg adjusted p-value threshold of 0.01 were considered for further selection.
2.) Exclude regions exhibiting a within-group standard deviation of average DMR methylation greater than 0.1 in any of the compared groups. For the neonatal vs postneonatal comparison, this cutoff was reduced to 0.05 due to the high number of adult HSC replicates.

DMRs passing this selection procedure were deemed genuine epigenetic scars. From the comparison between prenatal and postnatal we retained 100 epigenetic scars, from the comparison between neonatal and post-neonatal we retained 157.

### Linking transcriptional with epigenetic changes through generalized linear models

Generalized linear model with stepwise feature elimination using AIC was applied to identify DEGs associated with DMRs in a region of 200 Mb around each of the transcription start sites (TSS). The goal of this analysis was to identify genes whose alterations in expression could principally be attributed to changes in neighboring CpG methylation.

Specifically, our interest lay in genes differentially expressed between HM and non-HM cells derived from 10X data. Differentially expressed genes (DEGs) were identified using *Seurat* “findMarkers”, selecting genes meeting an adjusted p-value cutoff of 0.01. RNA expression data was normalized as outline in section: Pseudotemporal Analysis of Hematopoietic Lineages.

Likewise, DMRs were determined by comparing HM and non-HM patients utilizing bulk WGBS data. To minimize the influence of developmental regions, we excluded DMRs that overlapped with DMRs from all pairwise comparisons of non-tumor neonatal, juvenile, and adult patients. Overlap assessment was performed using the *IRanges* “findOverlaps” function (version 2.32.0) with default arguments. Subsequently, we computed the average methylation per patient per DMR.

After defining the sets of DEGs and DMRs, we calculated all DMRs that overlapped with a 100 kb region surrounding the gene body of each DEG. Prior to this, *hgLiftOver* (https://genome.ucsc.edu/cgi-bin/hgLiftOver) was employed to convert gene coordinates from hg38 to hg19.

Utilizing the “glm” function from the *stats* package in base R, we fitted a generalized linear model with a Gaussian link function to predict the expression of a given DEG through the methylation of associated DMRs. To reduce the number of model parameters (i.e., the number of DMRs predicting the expression of a given DEG), we employed the step function in base R, which optimizes the Akaike information criterion (AIC) in a stepwise manner. Briefly, this approach finds a balance between removing DMRs with minimal predictive power and maintaining acceptable model performance. Each DEG with at least one associated DMR after AIC optimization was considered linked to epigenetic changes.

Model performance was evaluated using standard metrics, including root-mean-square error, mean absolute error, Pearson correlation between prediction and ground truth, and ranked mean absolute error, which, unlike mean absolute error, utilizes the difference of ranks. DEGs were categorized based on the cellular location of their gene product using data from the *CellPhoneDB* database (https://raw.githubusercontent.com/Teichlab/cellphonedb-data/master/data/sources/protein_curated.csv). DEGs exhibiting a Pearson correlation between prediction and ground truth greater than 0.9 were selected as candidates for further analysis.

The heatmaps display expression and methylation values for all DMR-DEG associations for all candidate genes per patient and subgroup. Normalized gene expression was scaled to minimum and maximum across patients. Average methylation for each DMR is the average beta-values across all CpGs per region and JMML epitype. ‘Correlation’ depicts the Pearson correlation coefficients as per DMR-DEG association. The expression of 13 genes encoding for cell surface proteins was found to be significantly correlated with DNA methylation changes in up to 7 DMRs.

### *Ptpn11^E76K/+^* JMML mouse model

*Ptpn11^E76K/+^ Scl*-CreER^T^ mice were bred by crossing previously described *Scl*-CreER^T^ and *Ptpn11*^E76K-neo/+^ mice which were both on a B6.SJL-Ptprc^a^ Pepc^b^/BoyJ (CD45.1) genetic background^100,101^. All mice were kept at the DKFZ in individually ventilated cages under specific pathogen-free (SPF) conditions.

For all experiments, *Ptpn11^E76K/+^ Scl-*CreER^T^ and *Ptpn11^+/+^ Scl*-CreER^T^ mice at the age of 6 to 12 weeks were injected intraperitoneally (i.p.) on 5 consecutive days each day with 2 mg tamoxifen (dissolved in 200 µl sunflower oil with 10% ethanol). For differential blood count analysis, peripheral blood was collected from the vena facialis and measured using HemaVet950 (Drew Scientific) or scil Vet abc Plus+ (scil) veterinary hematology analyzers. Additionally, 30 µl of peripheral blood were used for flow cytometry as described below.

### Flow cytometric analysis of hematopoietic cells

Mice were sacrificed 18 weeks post injection by cervical dislocation according to European guidelines. Bone marrow of tibiae, femora, iliae and vertebrae were isolated by crushing the bones in IMDM (Gibco) three times. Additionally, one femur was flushed using 1 ml PBS supplemented with 2% FCS (PBS/FCS) to determine bone marrow cellularity and the cell suspension was afterwards added to the crushed bones. Spleen cells were isolated by passing the organs through a 40 µm cell strainer (Falcon). Subsequently, bone marrow cells, splenocytes and peripheral blood were resuspended in ACK lysis buffer (Lonza) for red blood cell lysis. Cells were washed and stained in PBS/FCS using established cell surface marker combinations for the identification of hematopoietic cell types (Supplementary Table 26). Flow cytometry analysis was performed using BD LSR II, BD LSR Fortessa and BD FACS Celesta (BD Biosciences).

### Bone marrow isolation and lineage depletion

Bone marrow cell isolation and lineage depletion have been performed as described before^102^. In short, bone marrow was isolated by crushing the bones three times in IMDM (Gibco) followed by red blood cell lysis using ACK lysis buffer (Lonza). Cells were washed in PBC/FCS and biotinylated antibodies added for lineage depletion (Supplementary Table 26). Mouse depletion Dynabeads (Invitrogen) were used for the depletion of lineage-labeled cells using a DynaMag-15 magnet (Invitrogen). Subsequently, lineage-negative cells were stained with antibodies for the isolation of HSPCs followed by FACS using BD FACS Aria II and III cell sorters (BD Biosciences).

### Single-cell genotyping

*Ptpn11^E76K/+^ Scl-*CreER^T^ mice were sacrificed three days after the last tamoxifen injection and subsequently lineage-depleted bone marrow cells isolated from tibiae, femora, iliae and vertebrae as described above. Single HSCs (Lin-cKIT+ Sca1+ CD48-CD150+) were FACS-isolated and expanded for 12 days in StemSpan SFEM (Stemcell Technologies) containing 1% Penicillin/Streptomycin (Gibco), 2 mM L-Glutamine (Gibco), 50 ng/ml IL-3 (PeproTech), 50 ng/ml IL-6 (PeproTech), 50 ng/ml TPO (PeproTech), 50 ng/ml mSCF (PeproTech) and 50 ng/ml Flt3L (PeproTech) at 37°C with 5% CO_2_ in 96-well U-bottom plates. Wells with colony growth were analyzed for induction of the *Ptpn11^E76K^* mutation by genotyping PCR^101^. In short, lysis buffer containing 0.01% Tween-20 (Sigma-Aldrich) and Proteinase K (Thermo Scientific) was added to the cells followed by incubation at 65°C for 60 min and 95°C for 15 min. Cell lysate was subjected to genotyping using a HotStar Taq polymerase (QIAGEN) and previously established primers^101^. PCR products were analyzed on a 2% agarose gel.

### Analysis of moribund mice

Moribund mice were sacrificed by cervical dislocation according to European guidelines. Spleen and live weight were determined and bone marrow cells isolated from tibiae, femora and iliae by flushing these bones with PBS/FCS. Flow cytometry was performed as described above using established cell surface marker combinations (Supplementary Table 26).

### Single-cell RNA-sequencing or material from *Ptpn11^E76K^* mice

Mice were sacrificed 18 weeks post injection by cervical dislocation according to European guidelines. Femur cellularity was determined by flushing femora with PBS/FCS. Additionally, bone marrow cells from vertebrae, femora, tibiae, iliae, humeri and sterna were isolated by crushing these bones three times in IMDM (Gibco). The femur cells were added to the crushed bones followed by red blood cell lysis using ACK lysis buffer (Lonza). Afterwards, 5 x 10^6^ cells were used for the staining of differentiated cells and 2.5 x 10^7^ cells for flow cytometry measurements (see ‘Flow cytometric analysis of hematopoietic cells’). The remaining cells were subjected to lineage depletion as described above. Lineage-negative and differentiated cells were stained with antibodies for the isolation of hematopoietic cell types and used for FACS (Supplementary Table 26). In total, 7,000 Lin-cKIT+ Sca1+ (LSK) and 7,000 Lin-cKIT+ (LK) cells were isolated from each mouse from lineage-negative bone marrow and 7,000 live total hematopoietic cells from unfractioned bone marrow. These cells were subjected to scRNA-sequencing using the Single Cell 3’ Reagent Kits v2 (10X Genomics) according to the manufacturer’s instructions. Sequencing was performed on a NovaSeq 6000 (Illumina) in paired-end (PE 26/96 bp) mode at the DKFZ GPCF.

Reads were aligned to GRCm38 and feature count matrices generated using Cell Ranger version 4.0.0 (10X Genomics). Feature counts were further processed using *Seurat* (version 4.0)^103^ whereby cells with more than 5% of mitochondrial reads or less than 500 and more than 5800 detected genes were removed from the analysis. Moreover, only genes that were expressed in a minimum of 10 cells were retained for further analysis. Doublets were identified and removed using *DoubletFinder*^104^. Cell cycle states were annotated based on the expression of stage specific marker genes (https://github.com/hbc/tinyatlas). Filtered data was subsequently integrated across genotypes using the “SCTransform” method followed by PCA, UMAP calculation and clustering. Clusters were annotated to hematopoietic cell states based on the expression of previously identified state-specific marker genes ^105,106^. Differentiation pseudotime was calculated using *slingshot* for the monocytic differentiation lineage and cells were ordered along this pseudotime based on their rank ^107^.

Cluster contributions were calculated based on 10,000 randomly selected cells from each genotype. P-values were determined using Fisher’s exact test followed by Benjamini-Hochberg’s false discovery rate (FDR) correction.

Transcriptional priming was calculated based on previously published gene sets ^108^ using the *Seurat* “AddModuleScore” function. Statistical significance was determined based on two LMMs as implemented in the *lme4* R package followed by ANOVA:

LMM1 = signature ∼ genotype + (1|mouse_id)

LMM2 = signature ∼ (1|mouse_id)

Murine fetal and adult HSC signature genes were previously published ^27^ and used for GSEA. In short, the *Seurat* “FindMarkers” function (logfc.threshold = -Inf, min.pct = 0.05, max.cells.per.ident = 1000) was used to calculate differential gene expression and subsequently genes not mapping to sex chromosomes were ranked based on fold change followed by GSEA using the *fgsea* package (https://www.biorxiv.org/content/10.1101/060012v3).

### Primary xenotransplantation

Newborn *Rag2^-/-^gc^-/-^* mice were used for xenotransplantation within the first four days of life. Mice were irradiated with 2,5 Gy. Six to eight hours after irradiation JMML (MNCs) were thawed, depleted from CD3^+^ T cells using CD3 MicroBeads (Miltenyi Biotec) and 1×10^6^ cells were injected intra-hepatically in 30 µl of PBS. Six to seven weeks after xenotransplantation, peripheral blood was obtained to inspect for human cell engraftment using flow cytometry with an antibody specifically detecting human pan-leukocyte antigen (hCD45). Successfully engrafted mice were then treated with alemtuzumab (100 µg/kg) or vehicle 1x/week for 4 weeks.

### *In vivo* anti-CD52 treatment

At eight weeks following xenotransplantation, mice were divided into two groups: the control group received 100 µl of PBS and the experimental group received 100 mg/kg anti-CD52 antibody (alemtuzumab, Genzyme Corporation Cambridge) prepared in 100 µl of PBS. Injections were given via tail vein once per week for four weeks.

One week after the end of the treatment, mice were sacrificed and blood, bone marrow, spleen, liver, and lungs were used for the analysis. Single cell suspensions were obtained for blood, bone marrow, and spleen, while liver and lungs were firstly digested with collagenase D (1 mg/ml) and DNase (25 mg/ml) for 30 minutes at 37°C followed by density gradient centrifugation. Cell suspensions were exposed to red blood cell lysis and stained with antibodies (Supplementary Table 27). All antibodies were used in 1:100 ratio. Cell acquisition and measurement was done using BD Fortessa. Different cell populations were characterized following gating strategy shown in Extended Data Figure 12i.

The expression levels are quantified as total number of events of engrafted CD45+ cells or absolute number of living cells calculated out from two femurs. Statistical analyses were performed using the unpaired Mann-Whitney test in GraphPad Prism 10 software. P values less than 0.05 were considered statistically significant. Treatment effect of alemtuzumab was calculated as difference of the percentage of engrafted human CD45 cells in treated vs untreated mice (treatment effect=treated_mean_ - untreated_mean_).

### Serial transplantations

For serial transplantations, 5-weeks old mice were sublethally irradiated with 3 Gy and six to eight hours after irradiation 1×10^7^ total bone marrow cells control- or alemtuzumab-treated mice were injected. Six weeks after transplantation, human cell engraftment was assessed in murine peripheral blood by flow cytometry using an antibody specifically detecting human pan-leukocyte antigen (hCD45). Terminally sick mice were eliminated and organs were analyzed as described above.

### Statistical analysis and data visualization

Statistical tests are specified in the corresponding figure legends and methods sections. If not differently specified, default statistical tests implemented in *Seurat* or *DSS* were used for scRNA-seq or WGBS analyses, respectively. For the analysis of JMML PDX mice, GraphPad Prism was used. Flow cytometry data was analyzed using FlowJo (v.10.6.2, v.10.8.1, BD Biosciences). Flow cytometers and sorters were operated using DIVA v.8 or v.9. Visualizations were generated using *ggplot2* and *pheatmap, Seurat* and *Nebulosa* in R version 4.0.3, 4.0.5, and 4.2.0^82^. For Kaplan-Meier analysis, the *survminer* R package was used to calculate leukemia-free survival (https://github.com/kassambara/survminer).

## Data availability

| Data | Source | Reference |
| --- | --- | --- |
| JMML_scRNA-seq | EGAS00001007357 | this study |
| FET_scRNA-seq | Human Cell Atlas | Popescu et al. (2019)<sup>68</sup> |
| NEO_scRNA-seq | Human Cell Atlas | https://data.humancellatlas.org/explore/projects/cc95ff89-2e68-4a08-a234-480eca21ce79 |
| JUV_scRNA-seq | HTA4_31 & HTA4_32 | this study |
| ADU_scRNA-seq | Human Cell Atlas | Hay et al. (2018)<sup>66</sup> |
| JMML_EPIC | EGAS00001007357 | this study |
| JMML_WGBS | EGAS00001007357 | this study |
| FET_WGBS | EGAS00001007357 | this study |
| NEO_WGBS | EGAS00001007357 | Maag, Toth et al. (in preparation) |
| JUV_WGBS | EGAS00001007357 | this study |
| ADU_WGBS | EGAS00001007357 | this study |
| Ptpn11-E76K_scRNA-seq | GEO: GSE240460 | this study |

## Code availability

WGBS and scRNA-seq data were processed and analyzed using publicly available software packages as specified in the corresponding methods sections. R code is available at https://github.com/maurerv/jmml_oncofetal_reprogramming.

## Acknowledgements

We thank the patients and their families and the EWOG-MDS study group for supporting our research. We also thank the NGS Core Facility, the Omics IT and Data Management Core Facility, the Flow Cytometry Core Facility and the Center for Preclinical Research, German Cancer Research Center (DKFZ), for providing excellent services. Emely Kleinert and Oliver Mücke provided excellent technical support. The Ptpn11 E76K neo/+ mouse was kindly shared by Dr. Cheng-Kui Qu, Department of Pediatrics, Aflac Cancer and Blood Disorders Center, Emory University, Atlanta, GA, USA. We thank all the members of the Division of Translational Medical Oncology, the Division of Applied Functional Genomics, Oliver Stegle, Stephen J. Clark and Wolf Reik for many fruitful discussions. This work was in part funded by Wellcome Trust (personal fellowship to S.B., institutional grant to the Wellcome Sanger Institute; references 220540/Z/20/A and 223135/Z/21/Z), by DFG (Emmy Noether RA 3166/1) to S.R., by the DKFZ International PhD Program and the German-Israeli Helmholtz International Research School “Cancer-TRAX” to M.Hakobyan, by NCT Molecular Precision Oncology Program to S.F., and by grants from the “Deutsche José Carreras Leukämie-Stiftung” (DJCLS 14R/2022) to M.M. and D.B.L. and from the “Wilhelm Sander-Stiftung” (# 2022.010.1) to D.B.L. This submission is part of the 2023 HCA Publication Package (HCA-105).

## Contributions

▪ M.H., M.S. and D.B.L. conceived the study, designed experiments and coordinated the project.
▪ R.R., K.M.B., T.B., S.R., S.H., C.B., L.J., M.Haniffa., C.M.N., C.F. and M.E. collected patient and/or healthy reference samples.
▪ M.S. established the Ptpn11^E76K^ mouse model and performed mouse experiments under supervision of D.B.L.
▪ M.H., M.S., K.B., J.P.M., J.R., M.Hakobyan, S.S., J.L., K.M.B., F.A., L.J., S.B., D.V., A.H.M., S.H., D.L. performed experiments.
▪ M.H., M.S., V.M., L.H., J.H., E.K., K.T., C.C., V.F. performed bioinformatic analyses.
▪ M.J.B., J.H., M.Schlesner and D.B.L. supervised bioinformatic analyses.
▪ J.R. and M.E. designed and J.R. performed the PDX experiments.
▪ M.H., M.S., J.R., V.M., L.H., J.H., P.L., M.D.M., S.H., S.Behjati, M.J.B., S.F., C.M.N., C.F., C.P., M.E., M.Schlesner and D.B.L. interpreted the data.
▪ D.B.L. acquired funding and supervised the project.
▪ M.H., M.S., and D.B.L. wrote the manuscript.

All authors read, edited and approved the manuscript.

## Ethics declarations – competing interests

A patent application has been filed by M.H., M.E. and D.B.L. based on the data described in this study. D.B.L. works as a drug safety physician for Infectopharm. J.H. is working as a consultant for CytoReason.

All other authors declare no competing interests.

