## Supplementary Figures for "Oncogenic RAS-Pathway Activation Drives Oncofetal Reprogramming and Creates Therapeutic Vulnerabilities in Juvenile Myelomonocytic Leukemia"

**Extended data figures and tables**

Hartmann & Schönung et al. 2023

**Supplementary information**

Excel file ‘Supplementary_Tables” contains Supplementary Tables 1 to 27, including scRNA-seq datasets, WGBS datasets, flow cytometry samples, murine Ptpn11$^{E76K}$ scRNA-seq datasets, PDX samples, used transcription signatures from literature, and antibodies used for cell sorting or flow cytometry analyses. Extended Data Figures 1 to 12 can be found below.

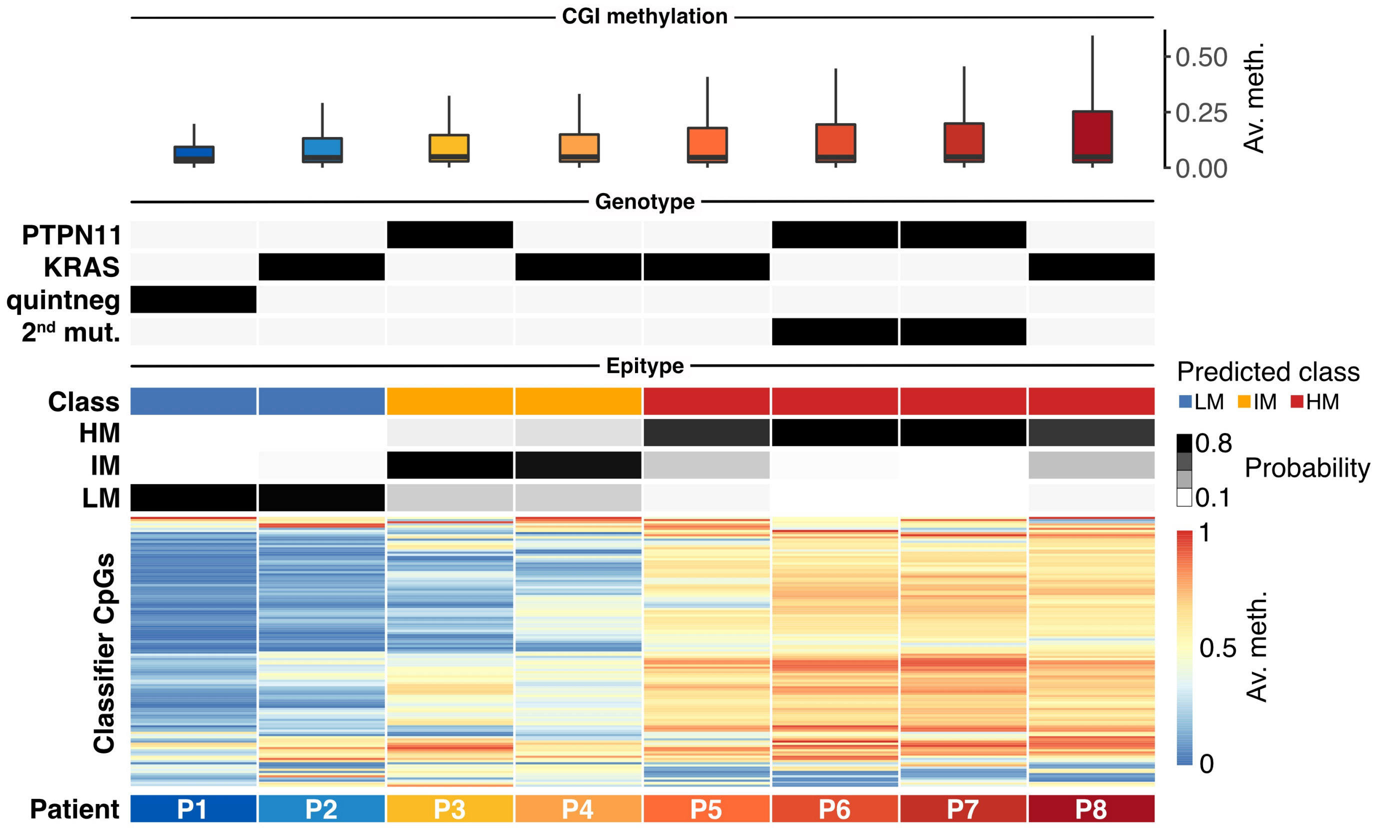

CGI methylation
0.50
0.25
0.00
Av. meth.
Genotype
PTPN11
KRAS
quintneg
2nd mut.
Epitype
Predicted class
LM IM HM
Class
HM
IM
LM
0.8
Probability
0.1
Classifier CpGs
1
0.5
0
Av. meth.
Patient
P1
P2
P3
P4
P5
P6
P7
P8

**Extended Data Figure 1: Multi-omics cohort: annotation of patient cohort to JMML DNA methylation subgroups ('epitypes').** Patient stratification was conducted using our published JMML DNA methylation classifier[1]. Heatmap shows average methylation values across 124 classifier CpGs. Based on the probability per JMML subgroup, patients were assigned to one of three subgroups, namely LM (low methylation, blue), IM (intermediate methylation, yellow), or HM (high methylation, red). The annotation tracks specify the genetic background of the patients, which was determined by JMML panel sequencing. As boxplots, average methylation of genome-wide CpG island (CGI) methylation values are summarized. Based on those values and in a semi-quantative manner, patients were annotated to a patient-specific color throughout the manuscript.

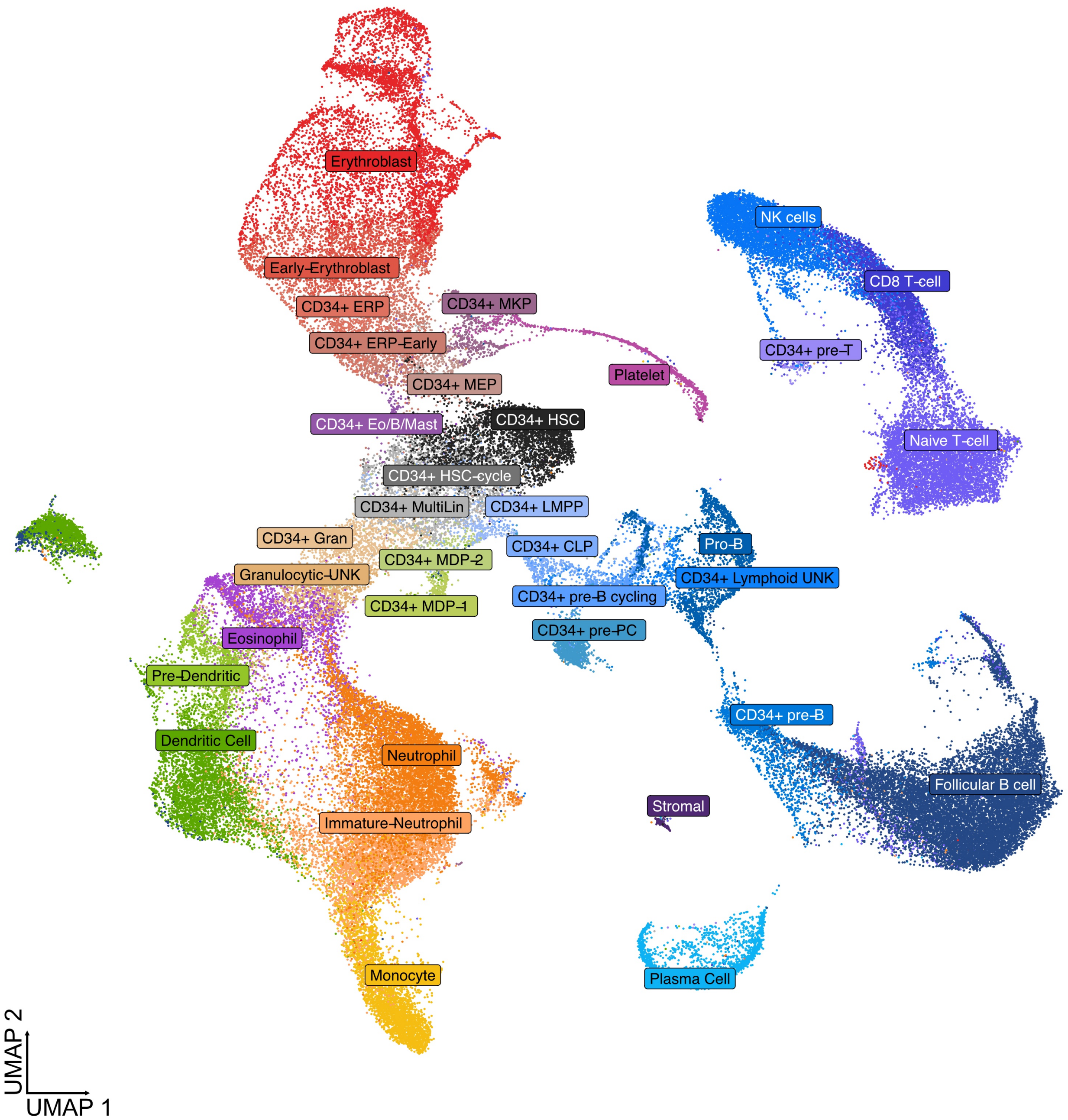
Erythroblast
Early-Erythroblast
CD34+ ERP
CD34+ MKP
CD34+ ERP-Early
Platelet
CD34+ MEP
CD34+ Eo/B/Mast
CD34+ HSC
CD34+ HSC-cycle
CD34+ MultiLin
CD34+ LMPP
CD34+ Gran
CD34+ CLP
CD34+ MDP-2
Granulocytic-UNK
CD34+ MDP-1
CD34+ pre-B cycling
CD34+ pre-PC
Eosinophil
Pre-Dendritic
Dendritic Cell
Neutrophil
Immature-Neutrophil
Monocyte
NK cells
CD8 T-cell
CD34+ pre-T
Naive T-cell
Pro-B
CD34+ Lymphoid UNK
CD34+ pre-B
Follicular B cell
Stromal
Plasma Cell
UMAP 2
UMAP 1

**Extended Data Figure 2**: **UMAP summarizing the scRNA-seq data of total hematopoiesis in JMML, neonatal cord blood, as well as juvenile and adult bone marrow.** Single-cells are colored by cell types, which were annotated by automated cell type annotation using Seurat's 'label transfer' function with adult bone marrow as a reference. JMML (n=7) and juvenile bone marrow (n=2) data were generated for this study. Neonatal cord blood and adult bone marrow data were taken from the Human Cell Atlas project[2]. The same colors are used for the corresponding cell types throughout the manuscript.

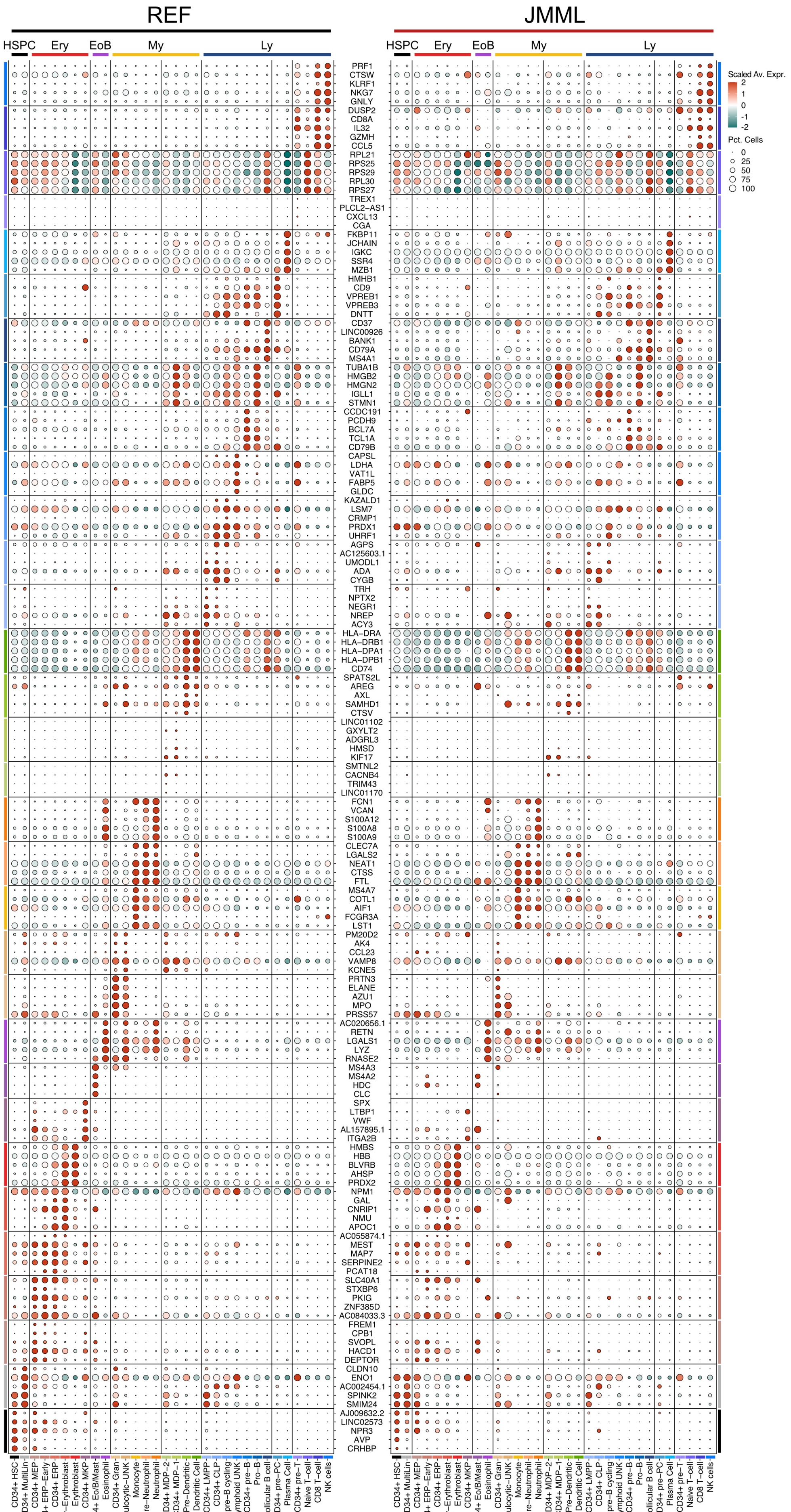

REF
JMML
HSPC
Ery
EoB
My
Ly
Scaled Av. Expr.
2
1
0
-1
-2
Pct. Cells
0
25
50
75
100
PRF1
CTSW
KLRF1
NKG7
GNLY
DUSP2
CD8A
IL32
GZMH
CCL5
RPL21
RPS25
RPS29
RPL30
RPS27
TREX1
PLCL2–AS1
CXCL13
CGA
FKBP11
JCHAIN
IGKC
SSR4
MZB1
HMHB1
CD9
VPREB1
VPREB3
DNTT
CD37
LINC00926
BANK1
CD79A
MS4A1
TUBA1B
HMGB2
HMGN2
IGLL1
STMN1
CCDC191
PCDH9
BCL7A
TCL1A
CD79B
CAPSL
LDHA
VAT1L
FABP5
GLDC
KAZALD1
LSM7
CRMP1
PRDX1
UHRF1
AGPS
AC125603.1
UMODL1
ADA
CYGB
TRH
NPTX2
NEGR1
NREP
ACY3
HLA–DRA
HLA–DRB1
HLA–DPA1
HLA–DPB1
CD74
SPATS2L
AREG
AXL
SAMHD1
CTSV
LINC01102
GXYLT2
ADGRL3
HMSD
KIF17
SMTNL2
CACNB4
TRIM43
LINC01170
FCN1
VCAN
S100A12
S100A8
S100A9
CLEC7A
LGALS2
NEAT1
CTSS
FTL
MS4A7
COTL1
AIF1
FCGR3A
LST1
PM20D2
AK4
CCL23
VAMP8
KCNE5
PRTN3
ELANE
AZU1
MPO
PRSS57
AC020656.1
RETN
LGALS1
LYZ
RNASE2
MS4A3
MS4A2
HDC
CLC
SPX
LTBP1
VWF
AL157895.1
ITGA2B
HMBS
HBB
BLVRB
AHSP
PRDX2
NPM1
GAL
CNRIP1
NMU
APOC1
AC055874.1
MEST
MAP7
SERPINE2
PCAT18
SLC40A1
STXBP6
PKIG
ZNF385D
AC084033.3
FREM1
CPB1
SVOPL
HACD1
DEPTOR
CLDN10
ENO1
AC002454.1
SPINK2
SMIM24
AJ009632.2
LINC02573
NPR3
AVP
CRHBP
CD34+ HSC
CD34+ MultiLin
CD34+ MEP
CD34+ ERP–Early
CD34+ ERP
Early–Erythroblast
Erythroblast
CD34+ MKP
CD34+ Eo/B/Mast
Eosinophil
CD34+ Gran
Granulocytic–UNK
Monocyte
Immature–Neutrophil
Neutrophil
CD34+ MDP–2
CD34+ MDP–1
Pre–Dendritic
Dendritic Cell
CD34+ LMPP
CD34+ CLP
CD34+ pre–B cycling
CD34+ Lymphoid UNK
CD34+ pre–B
Pro–B
Follicular B cell
CD34+ pre–PC
Plasma Cell
CD34+ pre–T
Naive T–cell
CD8 T–cell
NK cells

**Extended Data Figure 3**: **Dot plot summarizing the expression of the top 4 to 5 hematopoietic cell type marker genes as published in the human cell atlas project[2].** Average expression is scaled per gene. Dot sizes represent the percentage of cells per cell type expressing the corresponding gene. Expression values are separately analyzed for JMML and healthy references (‘REF’), including neonatal cord blood, as well as juvenile and adult bone marrow.

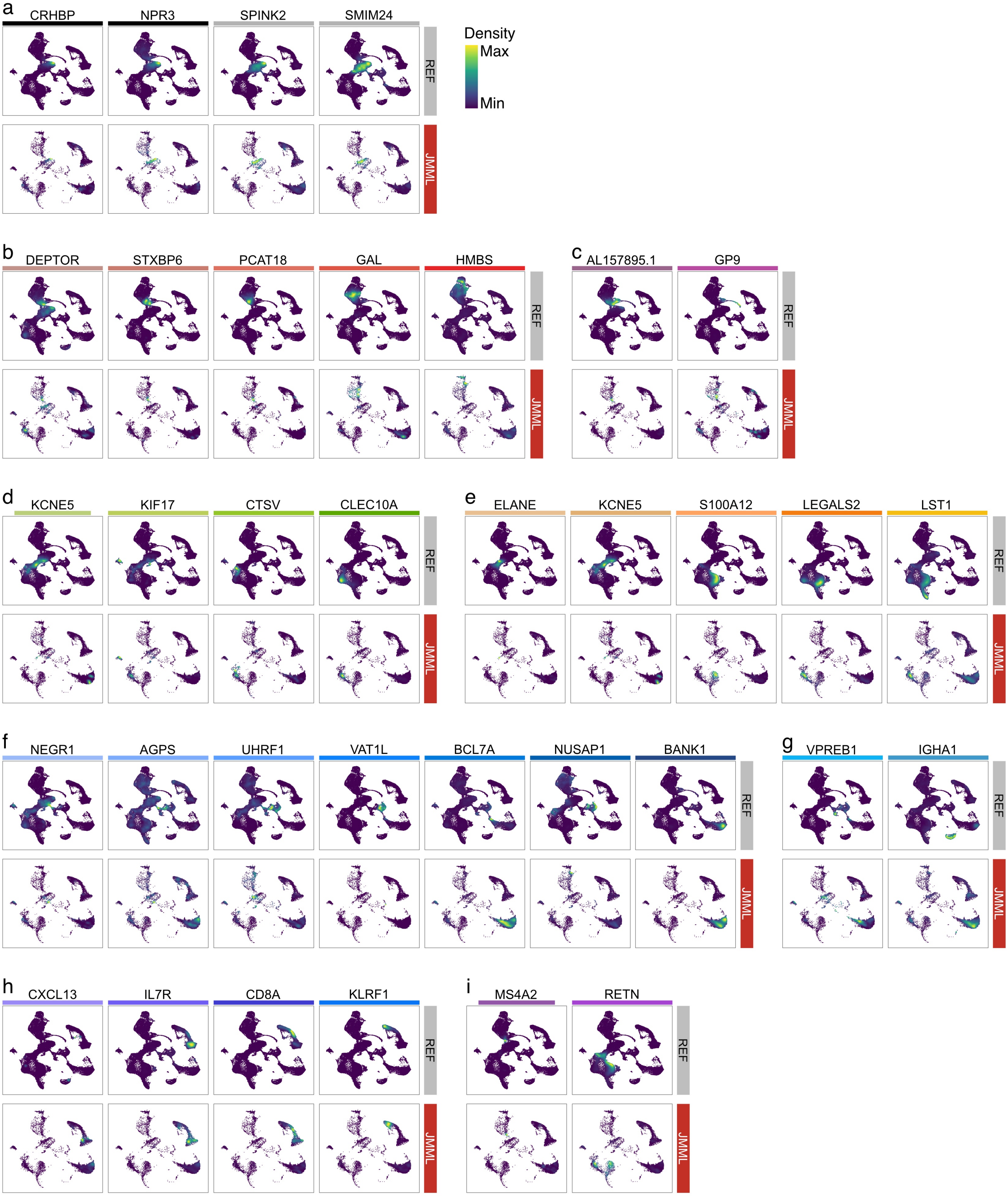

a
CRHBP
NPR3
SPINK2
SMIM24
Density
Max
Min
REF
JMML
b
DEPTOR
STXBP6
PCAT18
GAL
HMBS
REF
JMML
c
AL157895.1
GP9
REF
JMML
d
KCNE5
KIF17
CTSV
CLEC10A
REF
JMML
e
ELANE
KCNE5
S100A12
LEGALS2
LST1
REF
JMML
f
NEGR1
AGPS
UHRF1
VAT1L
BCL7A
NUSAP1
BANK1
REF
JMML
g
VPREB1
IGHA1
REF
JMML
h
CXCL13
IL7R
CD8A
KLRF1
REF
JMML
i
MS4A2
RETN
REF
JMML

**Extended Data Figure 4**: **Density plots depicting the cell type-specific expression of selected marker genes.** **a,** Hematopoietic stem and progenitor cells (HSPCs). **b,** Erythrocytes. **c,** Megakaryocytes. **d,** Dendritic cells. **e,** Monocytes and granulocytic neutrophils. **f,** B cells. **g,** Plasma cells. **h,** NK and T cells. **i,** Eosinophils.

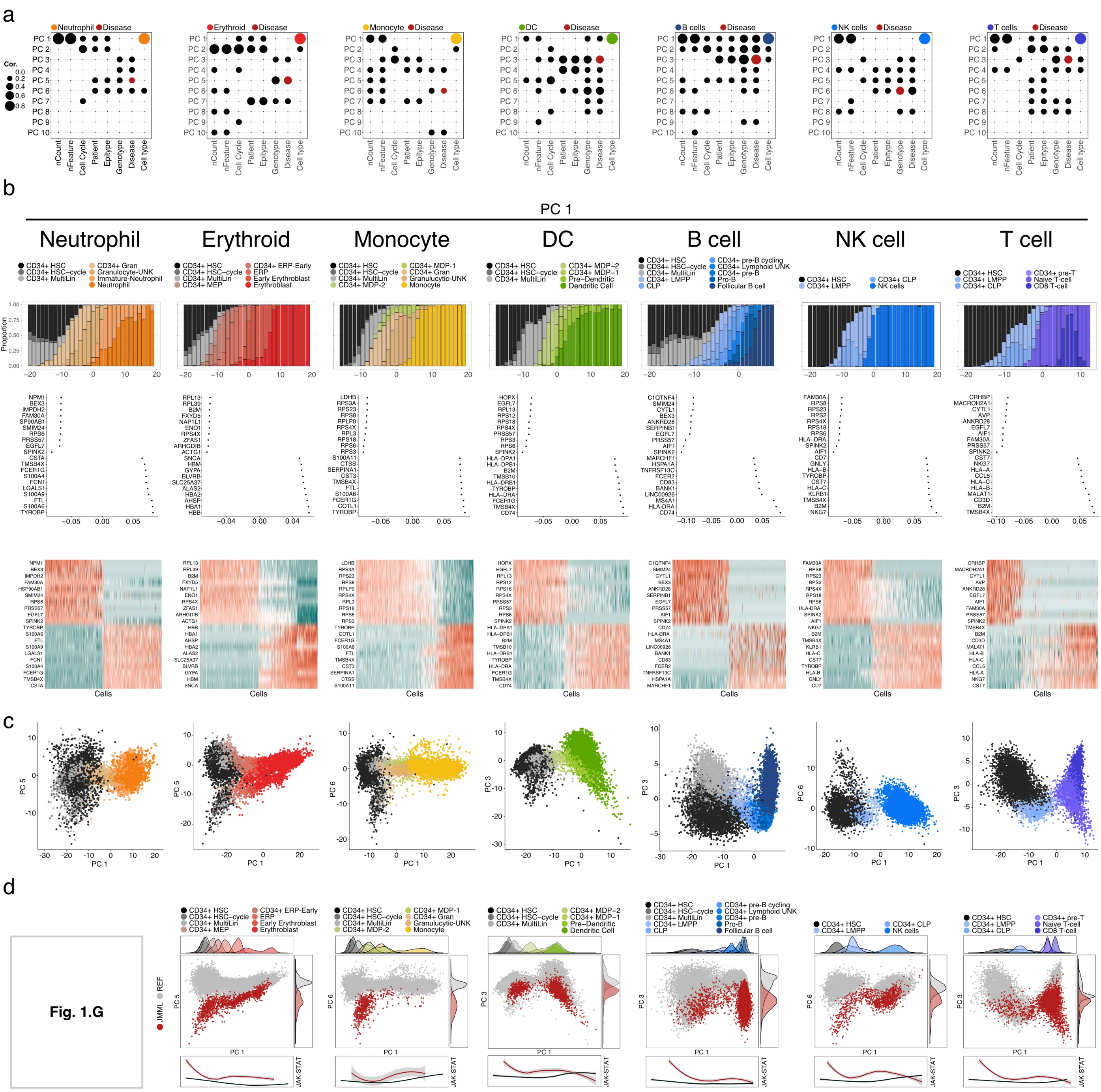
a
b
PC 1
Neutrophil
Erythroid
Monocyte
DC
B cell
NK cell
T cell
c
d
Fig. 1.G
JMML
REF
JAK-STAT

**Extended Data Figure 5**: **Principal component analyses (scRNA-seq data) of major hematopoietic lineages stratified by healthy references and JMML.** All analyses are ordered in columns per lineage as labeled in b. **a,** Dotplot of principal components. Depicted is the summary of the correlation of principal components with technical and biological features. nCount: number of transcripts/cell; nFeature: number of genes/cell; Cell cycle: cell cycle phase score; Patient: donor of primary material; Subgroup: JMML epitype; Genotype: type of RAS-pathway mutation; Disease: disease status, i.e. healthy or JMML; Cell type: annotated hematopoietic cell type. For every lineage, PC 1 was representative for differentiation (colored per lineage). Additionally, at least one PC was representative for disease-specific features (red) among the first six PCs. Those PCs were used for 2-D PCA plot visualizations (Figures 1g & ED5c,d). **b,** Detailed analysis of differentiation-specific PCs per lineage. From top to bottom: stacked bar plot of cell type contributions over the course of PC1; PC loadings and scaled normalized expression of top genes from PC1. **c,** PCA plot colored by cell types. **d,** PCA plot colored by disease state (JMML in red vs. healthy references in grey). Density plots of cell type distribution across PC1 or disease state for the second dimension. PCA plot for neutrophil lineage in Figure 1g.

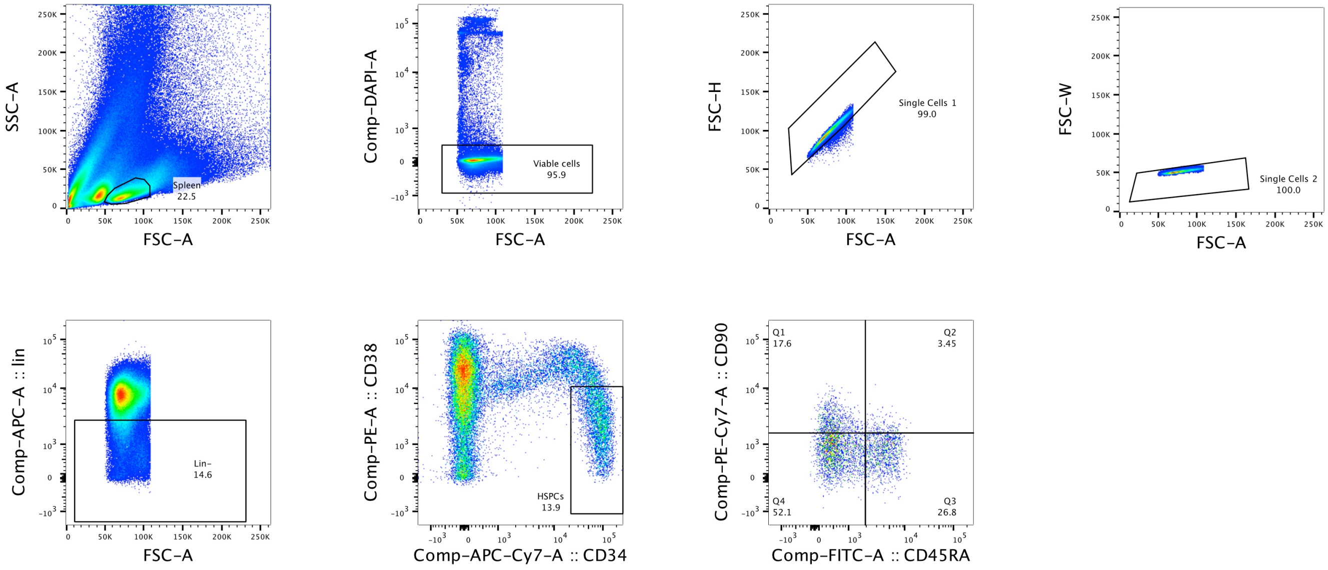
SSC-A
FSC-A
Spleen
22.5
Comp-DAPI-A
FSC-A
Viable cells
95.9
FSC-H
FSC-A
Single Cells 1
99.0
FSC-W
FSC-A
Single Cells 2
100.0
Comp-APC-A :: lin
FSC-A
Lin−
14.6
Comp-PE-A :: CD38
Comp-APC-Cy7-A :: CD34
HSPCs
13.9
Comp-PE-Cy7-A :: CD90
Comp-FITC-A :: CD45RA
Q1
17.6
Q2
3.45
Q3
26.8
Q4
52.1

**Extended Data Figure 6**: **Fluorescence activated cell sorting (FACS) strategy for in-depth analysis of JMML HSPCs.** $Lin^{-}CD34^{+}CD38^{lo/-}$ cells were selected for scRNA-seq analyses. $Lin^{-}CD34^{+}CD38^{lo/-}$CD45RA/CD90 quadrants were selected for ultra-low input WGBS analyses.

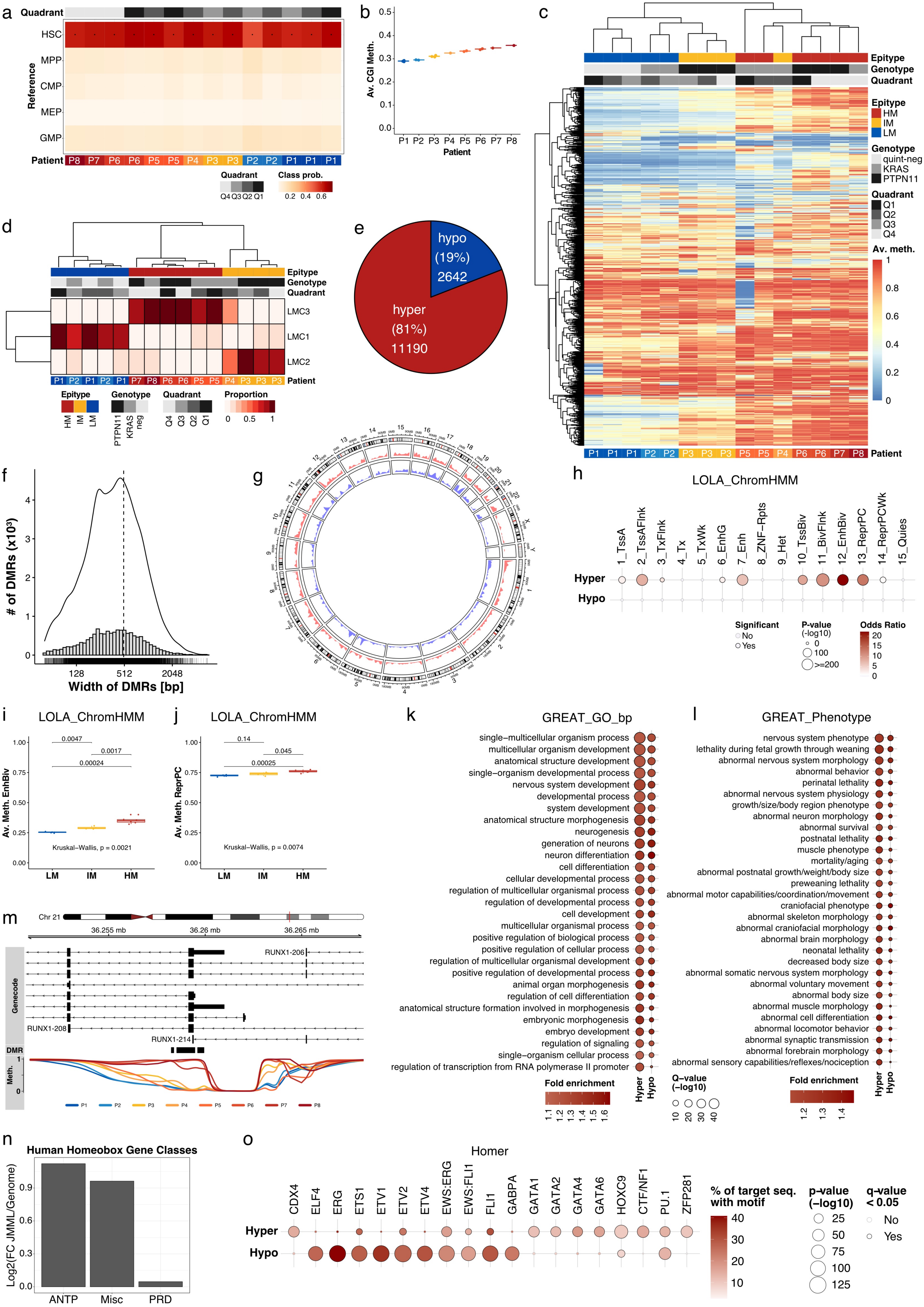
a
Quadrant
Reference
HSC
MPP
CMP
MEP
GMP
Patient
P8 P7 P6 P6 P5 P5 P4 P3 P3 P2 P2 P1 P1 P1
Quadrant
Q4 Q3 Q2 Q1
Class prob.
0.2 0.4 0.6
b
Av. CGI Meth.
Patient
P1 P2 P3 P4 P5 P6 P7 P8
c
Epitype
Genotype
Quadrant
HM
IM
LM
quint-neg
KRAS
PTPN11
Q1
Q2
Q3
Q4
Av. meth.
P1 P1 P1 P2 P2 P3 P3 P3 P5 P5 P4 P6 P6 P7 P8
Patient
d
Epitype
Genotype
Quadrant
LMC3
LMC1
LMC2
P1 P2 P1 P2 P1 P7 P8 P6 P6 P5 P5 P4 P3 P3 P3
Patient
Proportion
PTPN11
KRAS
neg
e
hypo
(19%)
2642
hyper
(81%)
11190
f
### of DMRs (x10³)
Width of DMRs [bp]
128
512
2048
g
h
LOLA_ChromHMM
1_TssA
2_TssAFlnk
3_TxFlnk
4_Tx
5_TxWk
6_EnhG
7_Enh
8_ZNF-Rpts
9_Het
10_TssBiv
11_BivFlnk
12_EnhBiv
13_ReprPC
14_ReprPCWk
15_Quies
Hyper
Hypo
Significant
No
Yes
P-value
(-log10)
100
>=200
Odds Ratio
i
LOLA_ChromHMM
Av. Meth. EnhBiv
0.0047
0.0017
0.00024
Kruskal–Wallis, p = 0.0021
LM
IM
HM
j
LOLA_ChromHMM
Av. Meth. ReprPC
0.14
0.045
0.00025
Kruskal–Wallis, p = 0.0074
k
GREAT_GO_bp
single–multicellular organism process
multicellular organism development
anatomical structure development
single–organism developmental process
nervous system development
developmental process
system development
anatomical structure morphogenesis
neurogenesis
generation of neurons
neuron differentiation
cell differentiation
cellular developmental process
regulation of multicellular organismal process
regulation of developmental process
cell development
multicellular organismal process
positive regulation of biological process
positive regulation of cellular process
regulation of multicellular organismal development
positive regulation of developmental process
animal organ morphogenesis
regulation of cell differentiation
anatomical structure formation involved in morphogenesis
embryonic morphogenesis
embryo development
regulation of signaling
single–organism cellular process
regulation of transcription from RNA polymerase II promoter
Fold enrichment
Hyper
Hypo
Q–value
(−log10)
l
GREAT_Phenotype
nervous system phenotype
lethality during fetal growth through weaning
abnormal nervous system morphology
abnormal behavior
perinatal lethality
abnormal nervous system physiology
growth/size/body region phenotype
abnormal neuron morphology
abnormal survival
postnatal lethality
muscle phenotype
mortality/aging
abnormal postnatal growth/weight/body size
preweaning lethality
abnormal motor capabilities/coordination/movement
craniofacial phenotype
abnormal skeleton morphology
abnormal craniofacial morphology
abnormal brain morphology
neonatal lethality
decreased body size
abnormal somatic nervous system morphology
abnormal voluntary movement
abnormal body size
abnormal muscle morphology
abnormal cell differentiation
abnormal locomotor behavior
abnormal synaptic transmission
abnormal forebrain morphology
abnormal sensory capabilities/reflexes/nociception
Fold enrichment
Hyper
Hypo
m
Chr 21
36.255 mb
36.26 mb
36.265 mb
RUNX1-206
Genecode
RUNX1-208
RUNX1-214
DMR
Meth.
P1 P2 P3 P4 P5 P6 P7 P8
n
Human Homeobox Gene Classes
Log2(FC JMML/Genome)
ANTP
Misc
PRD
o
Homer
CDX4
ELF4
ERG
ETS1
ETV1
ETV2
ETV4
EWS:ERG
EWS:FLI1
FLI1
GABPA
GATA1
GATA2
GATA4
GATA6
HOXC9
CTF/NF1
PU.1
ZFP281
Hyper
Hypo
% of target seq.
with motif
p–value
(−log10)
q–value
< 0.05
No
Yes

**Extended Data Figure 7**: **Ultra-low input whole-genome bisulfite sequencing data analysis of Lin$^-$CD34$^+$CD38$^{lo/-}$ HSPCs from primary JMML patient samples. a,** DNA methylation-based cell type classifier to determine the cell type-specific DNA methylation profiles across Lin$^-$CD34$^+$CD38$^{lo/-}$CD45RA/CD90 quadrants[3]. Normal bone marrow from adult donors was taken as a reference. HSC, Lin$^-$CD34$^+$CD38$^-$CD45RA$^-$CD90$^+$; MPP, Lin$^-$CD34$^+$CD38$^-$CD45RA$^-$CD90$^-$; CMP, Lin$^-$CD34$^+$CD38$^+$CD10$^-$CD45RA$^-$CD135$^+$; GMP, Lin$^-$CD34$^+$CD38$^+$CD10$^-$CD45RA$^+$CD135$^+$; MEP, Lin$^-$CD34$^+$CD38$^+$CD10$^-$CD45RA$^-$CD135$^-$. **b,** Boxplot of the average genome-wide DNA methylation of CpG islands (CGI) across patients. **c,** Heatmap of the average methylation of the 1,000 most variable promoters. Epitype, JMML DNA methylation subgroup; Genotype, RAS mutation; Celltype, Lin$^-$CD34$^+$CD38$^{lo/-}$ CD45RA/CD90 quadrant. **d,** Reference-free decomposition of DNA methylation data using MeDeCom[4] with the following settings: k = 3, lambda = 0.01. **e-g,** HM vs non-HM DMR calling. **e,** Piechart stratifying DMRs into hyper- (11,190) and hypo-methylated (2,642). **f,** Histogram summarizing the size distribution of JMML DMRs with a median of 512 bp. **g,** Circos plot with histograms summarizing genomic DMR locations. Red represents hyper- and blue represents hypomethylated regions in HM vs. non-HM DMRs. Genome-wide distribution of JMML DMRs. **h-j,** Locus overlap analysis for enrichment of genomic ranges (LOLA) using the 15-state ChromHMM chromatin feature annotation. **h,** Bubble plot summarizing genome-wide enrichment analysis of DMRs in chromatin features (ChromHMM) using locus overlap analysis (LOLA). **i,** Boxplot of average methylation in polycomb-repressed (ReprPC) domains across JMML epitypes. **j,** Boxplot of average methylation in bivalent enhancers (EnhBiv) across JMML epitypes. **k,l,** Genomic Regions Enrichment of Annotations Tool (GREAT)[5] analysis for gene ontology analysis of (K.) biological processes (GO_bp) and (L.) murine phenotypes. **m,** Locus line plot of DNA methylation at the RUNX1 locus across JMML patients. Smoothed DNA methylation; 'DMR' indicates HM vs. non-HM JMML DMRs. **n,** Bar plot depicting the logarithmic fold change of the number of homeobox genes associated to HM vs. non-HM DMRs relative to the genome-wide abundance. Homeobox gene classes were stratified as previously published[6]: ANTP, *Antennapedia* (incl. HOXL and NKL genes); PRD, *paired* (incl. PAX and PAXL genes); Misc incl. LIM, POU, HNF, SINE, CUT, PROS, ZF, CERS class genes. **o,** Bubble plot of the genome-wide analysis of enriched motifs across DMRs using Hypergeometric Optimization of Motif EnRichment (HOMER) to summarize the enrichment of TF binding motifs across HM vs non-HM DMRs.

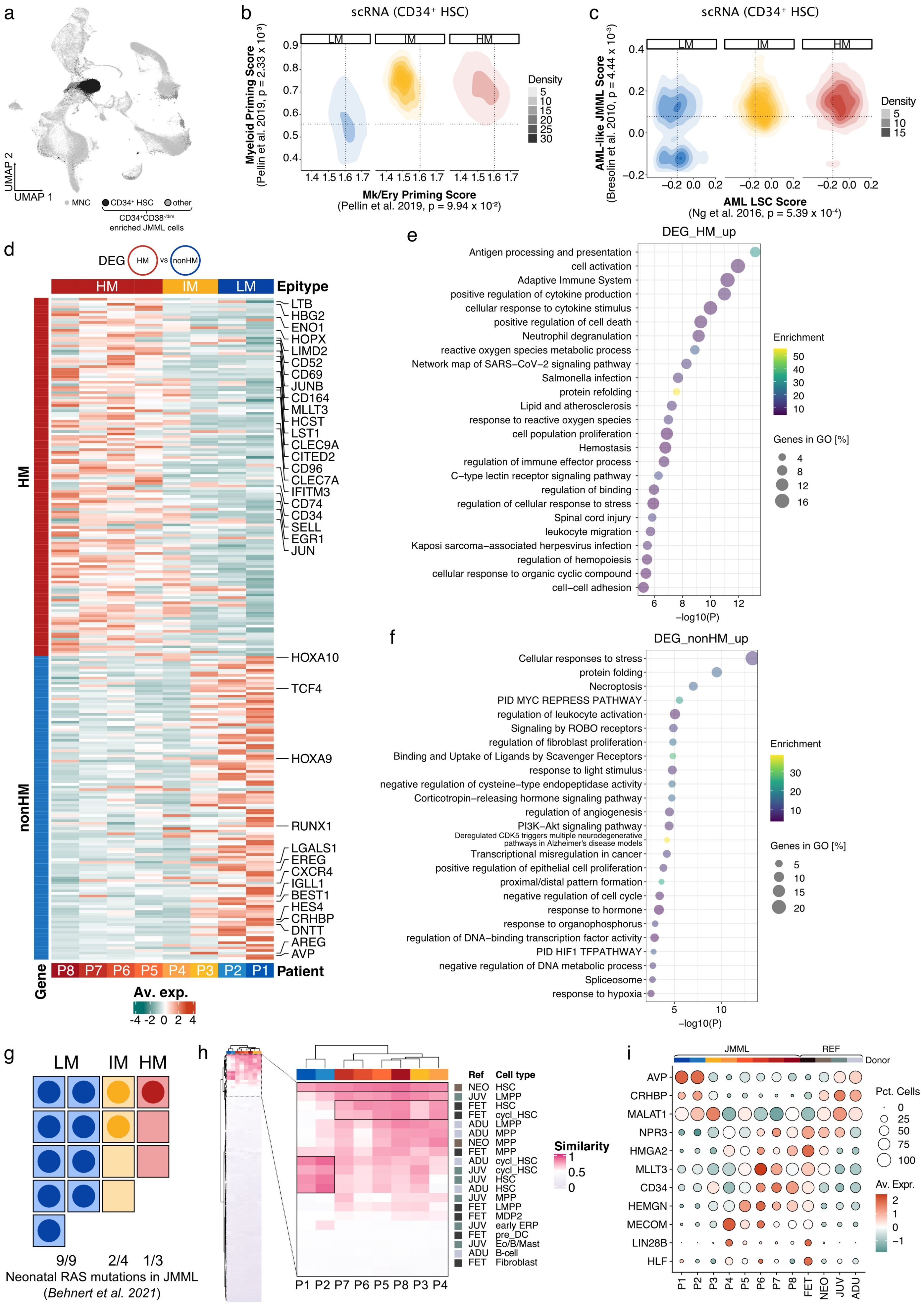

a
UMAP 2
UMAP 1
MNC
CD34⁺ HSC
other
CD34⁺CD38⁻/dim enriched JMML cells
b
scRNA (CD34⁺ HSC)
LM
IM
HM
Myeloid Priming Score
(Pellin et al. 2019, p = 2.33 x 10⁻³)
Mk/Ery Priming Score
(Pellin et al. 2019, p = 9.94 x 10⁻²)
Density
c
scRNA (CD34⁺ HSC)
AML-like JMML Score
(Bresolin et al. 2010, p = 4.44 x 10⁻³)
AML LSC Score
(Ng et al. 2016, p = 5.39 x 10⁻⁴)
Density
d
DEG HM vs nonHM
Epitype
HM
nonHM
LTB
HBG2
ENO1
HOPX
LIMD2
CD52
CD69
JUNB
CD164
MLLT3
HCST
LST1
CLEC9A
CITED2
CD96
CLEC7A
IFITM3
CD74
CD34
SELL
EGR1
JUN
HOXA10
TCF4
HOXA9
RUNX1
LGALS1
EREG
CXCR4
IGLL1
BEST1
HES4
CRHBP
DNTT
AREG
AVP
Gene
P8 P7 P6 P5 P4 P3 P2 P1
Patient
Av. exp.
e
DEG_HM_up
Antigen processing and presentation
cell activation
Adaptive Immune System
positive regulation of cytokine production
cellular response to cytokine stimulus
positive regulation of cell death
Neutrophil degranulation
reactive oxygen species metabolic process
Network map of SARS−CoV−2 signaling pathway
Salmonella infection
protein refolding
Lipid and atherosclerosis
response to reactive oxygen species
cell population proliferation
Hemostasis
regulation of immune effector process
C−type lectin receptor signaling pathway
regulation of binding
regulation of cellular response to stress
Spinal cord injury
leukocyte migration
Kaposi sarcoma−associated herpesvirus infection
regulation of hemopoiesis
cellular response to organic cyclic compound
cell−cell adhesion
−log10(P)
Enrichment
Genes in GO [%]
f
DEG_nonHM_up
Cellular responses to stress
protein folding
Necroptosis
PID MYC REPRESS PATHWAY
regulation of leukocyte activation
Signaling by ROBO receptors
regulation of fibroblast proliferation
Binding and Uptake of Ligands by Scavenger Receptors
response to light stimulus
negative regulation of cysteine−type endopeptidase activity
Corticotropin−releasing hormone signaling pathway
regulation of angiogenesis
PI3K−Akt signaling pathway
Deregulated CDK5 triggers multiple neurodegenerative pathways in Alzheimer's disease models
Transcriptional misregulation in cancer
positive regulation of epithelial cell proliferation
proximal/distal pattern formation
negative regulation of cell cycle
response to hormone
response to organophosphorus
regulation of DNA−binding transcription factor activity
PID HIF1 TFPATHWAY
negative regulation of DNA metabolic process
Spliceosome
response to hypoxia
−log10(P)
Enrichment
Genes in GO [%]
g
LM IM HM
9/9 2/4 1/3
Neonatal RAS mutations in JMML
(Behnert et al. 2021)
h
Ref Cell type
NEO HSC
JUV LMPP
FET HSC
FET cycl_HSC
ADU LMPP
ADU MPP
NEO MPP
FET MPP
ADU cycl_HSC
JUV cycl_HSC
JUV HSC
ADU HSC
JUV MPP
FET LMPP
FET MDP2
JUV early ERP
FET pre_DC
JUV Eo/B/Mast
ADU B-cell
FET Fibroblast
Similarity
P1 P2 P7 P6 P5 P8 P3 P4
i
JMML
REF
Donor
AVP
CRHBP
MALAT1
NPR3
HMGA2
MLLT3
CD34
HEMGN
MECOM
LIN28B
HLF
Pct. Cells
Av. Expr.
P1 P2 P3 P4 P5 P6 P7 P8 FET NEO JUV ADU

**Extended Data Figure 8**: **scRNA-seq data analysis of Lin$^-$CD34$^+$CD38$^{lo/-}$ HSPCs from primary JMML patient samples.** **a,** Projection of scRNA-seq data of 20,163 Lin$^-$CD34$^+$CD38$^{lo/-}$ enriched HSPCs onto the UMAP from Figure 1.B (total hematopoiesis, small light grey dots). Additionally sequenced JMML HSPCs are represented as larger dots (CD34$^+$ HSC in black and others in grey). **b,c,** Density plots of single-cell transcription scores of transcriptionally defined JMML HSCs (CD34$^+$ HSC) stratified by JMML epitype. **b,** Megakaryocytic/erythroid (Mk/Ery) vs. lympho/myeloid priming. These gene signatures reflect the first branching point in lineage commitment of human HSCs according to Pellin et al. (2019)[7]. **c,** Transcription scores for clinically aggressive JMML (AML-like JMML) as defined by Bresolin et al. (2010)[8] and leukemia stem cell programs (17-gene LSC) as defined by Ng et al. (2016)[9]. **d,** Heatmap of 255 differentially expressed genes (DEGs) identified between CD34$^+$ HSCs from HM and non-HM JMML patients. Depicted are normalized average expression levels aggregated across all HSCs within each of the patients. **e,f,** Bubble plots summarizing the top 25 significant terms of functional enrichment analysis of HM vs non-HM DEGs using Metascape[10]. Data was stratified by upregulated genes in HM or non-HM HSCs, respectively. Bubble size represents percentage of genes in GO and color the enrichment per term. **g,** Re-analysis of neonatal blood spot genotyping by Behnert et al. (2022)[11]. The frequency of RAS driver mutations in neonatal blood spots of individuals that developed JMML later in life is stratified by JMML epitype. Each square represents a single patient, dark dots represent RAS-positive blood spots and unfilled squares RAS-negative cases. **h,** Reference-based cell type similarity analysis (logistic regression) of JMML HSCs to a reference map of 116 hematopoietic reference cell types across developmental stages (rows). Values above 0.5 (pink) and below 0.5 (grey) refer to similarity and dissimilarity, respectively, to the reference cell type; 0.5 represents randomness (white). Left panel shows the entire results. Right panel shows a zoom into the top part or the heatmap which includes the references with positive similarity values. Row labeling refers to the reference stage (Ref: FET, fetal; NEO, neonatal; JUV, juvenile; ADU, adult) and the cell type, respectively. Reference cell types shown in the main figure 3b are marked by black boxes. **i.** Dot plot of manually curated HSC marker from pre- and postnatal references. The gene list is manually curated based on previously published data[2,7,12,13]. Average expression is scaled per gene. Dot sizes represent the percentage of cells expressing the corresponding gene.

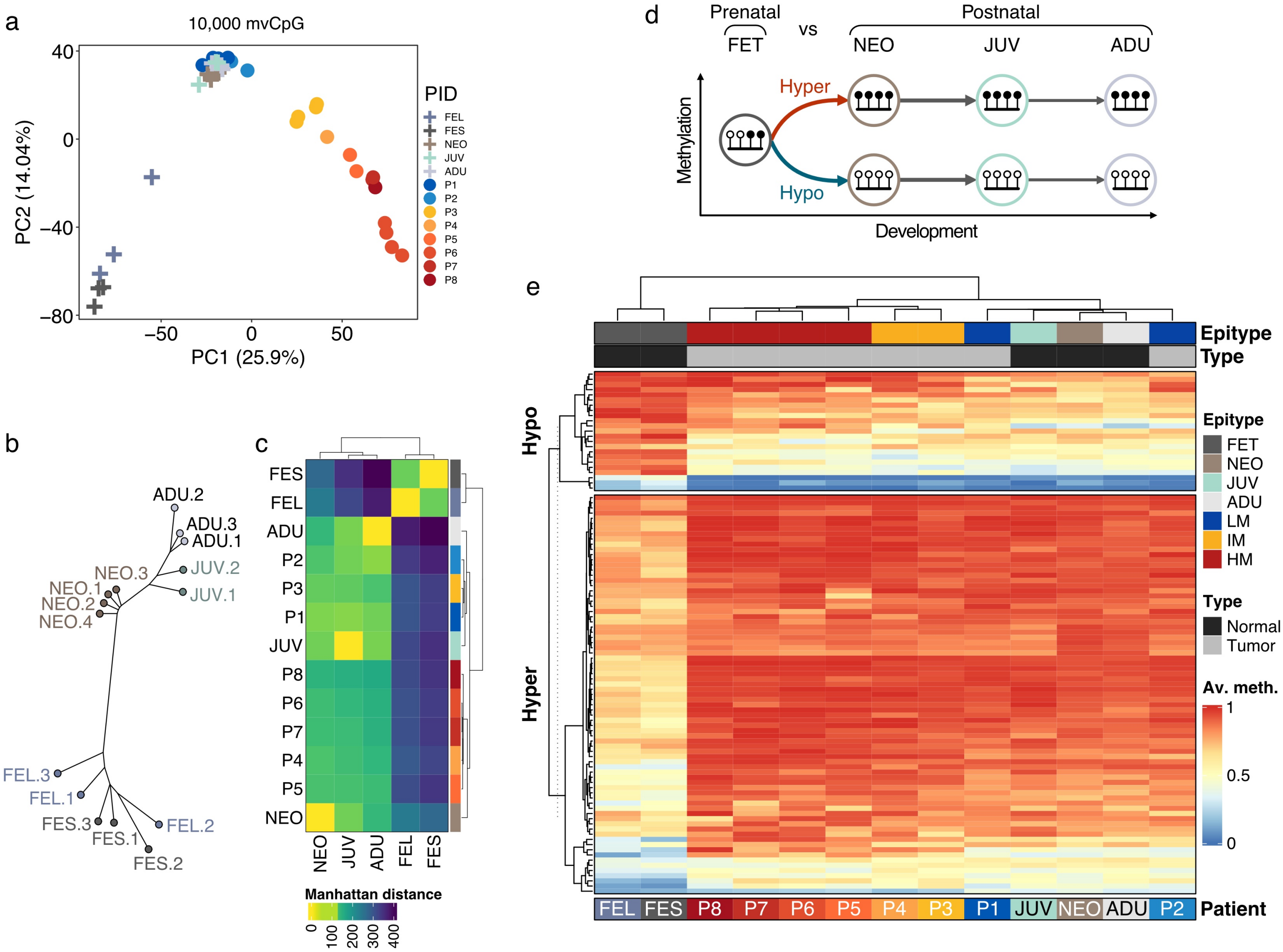

a
10,000 mvCpG
PC1 (25.9%)
PC2 (14.04%)
PID
FEL
FES
NEO
JUV
ADU
P1
P2
P3
P4
P5
P6
P7
P8
b
ADU.2
ADU.3
ADU.1
JUV.2
JUV.1
NEO.3
NEO.1
NEO.2
NEO.4
FEL.3
FEL.1
FEL.2
FES.3
FES.1
FES.2
c
Manhattan distance
d
Prenatal
FET
vs
Postnatal
NEO
JUV
ADU
Hyper
Hypo
Methylation
Development
e
Epitype
Type
Hypo
Hyper
FET
NEO
JUV
ADU
LM
IM
HM
Normal
Tumor
Av. meth.
Patient
FEL FES P8 P7 P6 P5 P4 P3 P1 JUV NEO ADU P2

**Extended Data Figure 9**: **Ultra-low input whole-genome bisulfite sequencing data analysis of Lin$^{-}$CD34$^{+}$CD38$^{lo/-}$ HSPCs from primary JMML patient samples in the context of normal HSC development.** Ultra-low input WGBS analysis of normal and JMML HSCs from primary samples. FET corresponds to both fetal spleen (FES) and fetal liver (FEL) samples. NEO is neonatal blood. JUV and ADU are juvenile and adult bone marrow. **a,** Principal component analysis (PCA) of the 10k most-variable CpGs (mvCpGs) across the entire WGBS dataset including reference and JMML HSCs. Low-risk HSCs cluster with postnatal HSCs. **b,c,** Methylation-based phylogenetic relationships between JMML and healthy reference HSCs across development using WGBS data. **b.** Phylogenetic tree analysis of healthy HSCs based on genomic regions defined in 3.F. Selected methylation dynamics reflect the expected order of normal HSC development. **c,** Heatmap showing the distances of samples in the phylogenetic tree. Manhatten distances were calculated based on the branch length per subgroup. **d,e,** 'Epigenetic scars' across normal and malignant HSCs. Epigenetic scars are per definition DNA methylation changes that occur at a single step during development but do not change downstream of this event. 'Hypo' and 'hyper' correspond to scars whose methylation de- or increase during developmental state transition. **d,** Schematic of epigenetic scars of the transition between fetal and postnatal stages **e,** Heatmap showing methylation beta values of epigenetic scars from the transition of fetal to postnatal HSCs. Hypo: loss of DNA methylation; hyper: gain of DNA methylation.

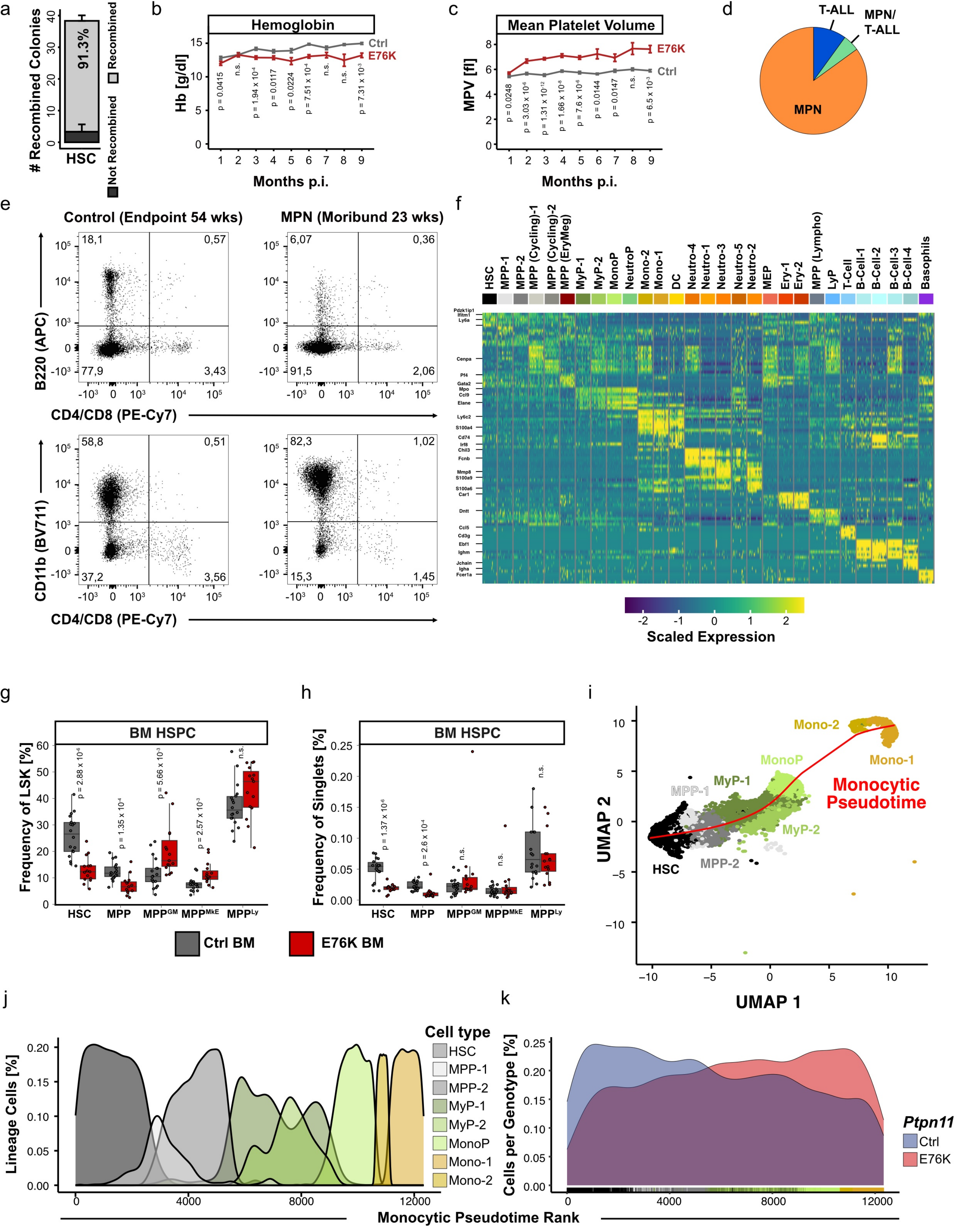

a
### Recombined Colonies
91.3%
HSC
Recombined
Not Recombined
b
Hemoglobin
Hb [g/dl]
Ctrl
E76K
Months p.i.
c
Mean Platelet Volume
MPV [fl]
E76K
Ctrl
Months p.i.
d
T-ALL
MPN/
T-ALL
MPN
e
Control (Endpoint 54 wks)
MPN (Moribund 23 wks)
B220 (APC)
CD4/CD8 (PE-Cy7)
CD11b (BV711)
f
Scaled Expression
g
BM HSPC
Frequency of LSK [%]
h
BM HSPC
Frequency of Singlets [%]
Ctrl BM
E76K BM
i
Monocytic Pseudotime
UMAP 1
UMAP 2
j
Cell type
Lineage Cells [%]
Monocytic Pseudotime Rank
k
Cells per Genotype [%]
Ptpn11
Ctrl
E76K

**Extended Data Figure 10: HSC-specific *Ptpn11*$^{E76K/+}$ JMML mouse model recapitulates human disease. a**, Recombination efficacy of the loxP-flanked PGK-Neo cassette in HSCs. *Ptpn11*$^{E76K/+}$ *Scl*-CreER$^{T}$ mice were injected at the age of 6 weeks on 5 consecutive days with tamoxifen and recombination of the loxP-flanked PGK-Neo cassette was analyzed after 72h in HSCs by PCR. **b-c**, Differential blood counts in *Ptpn11*$^{E76K/+}$ *Scl*-CreER$^{T}$ mice (E76K, n = 29) and *Ptpn11*$^{+/+}$ *Scl*-CreER$^{T}$ controls (Ctrl, n = 23). The results for hemoglobin levels (Hb; **b**) and mean platelet volume (MPV; **c**) are shown over a time course of 9 months. Mean and SEM are depicted. Statistical significances was determined using Student's t-test. **d**, Disease phenotype in moribund E76K mice (n = 20). **e**, Flow cytometry measurement of bone marrow cells from a control mouse and a E76K mouse with MPN. The percentage of cells in each gate is depicted. Axis labels indicate the respective surface markers and fluorophores. **f**, Expression of hematopoietic marker genes in scRNA-seq. The scaled expression of the top 5 marker genes for each of the 29 clusters identified in the scRNA-seq data is depicted as a heatmap. Each cluster was downsampled to 25 cells for visualization. Cluster-specific marker genes were empirically selected and labeled. **g-h**, Hematopoietic stem and progenitor compartment remodeling in E76K (n = 15) and Ctrl (n = 18) mice 18-weeks after tamoxifen induction. HSPC frequencies were determined using flow cytometry and are plotted as frequencies of LSK cells (**g**) and singlets (**h**). Whiskers are defined as 1.5 times interquartile range. Statistical significance was calculated using Student's t-test. **i**, UMAP embedding showing cells from the monocytic differentiation lineage and the calculated monocytic pseudotime. **j-k,** Density plots display the distribution of 3000 randomly selected bone marrow cells from E76K and Ctrl mice along the monocytic pseudotime. The distribution is colored by cell type (**j**) or genotype (**k**), respectively. The annotated cell types are visualized as bars below the density curve per genotype (**k**).

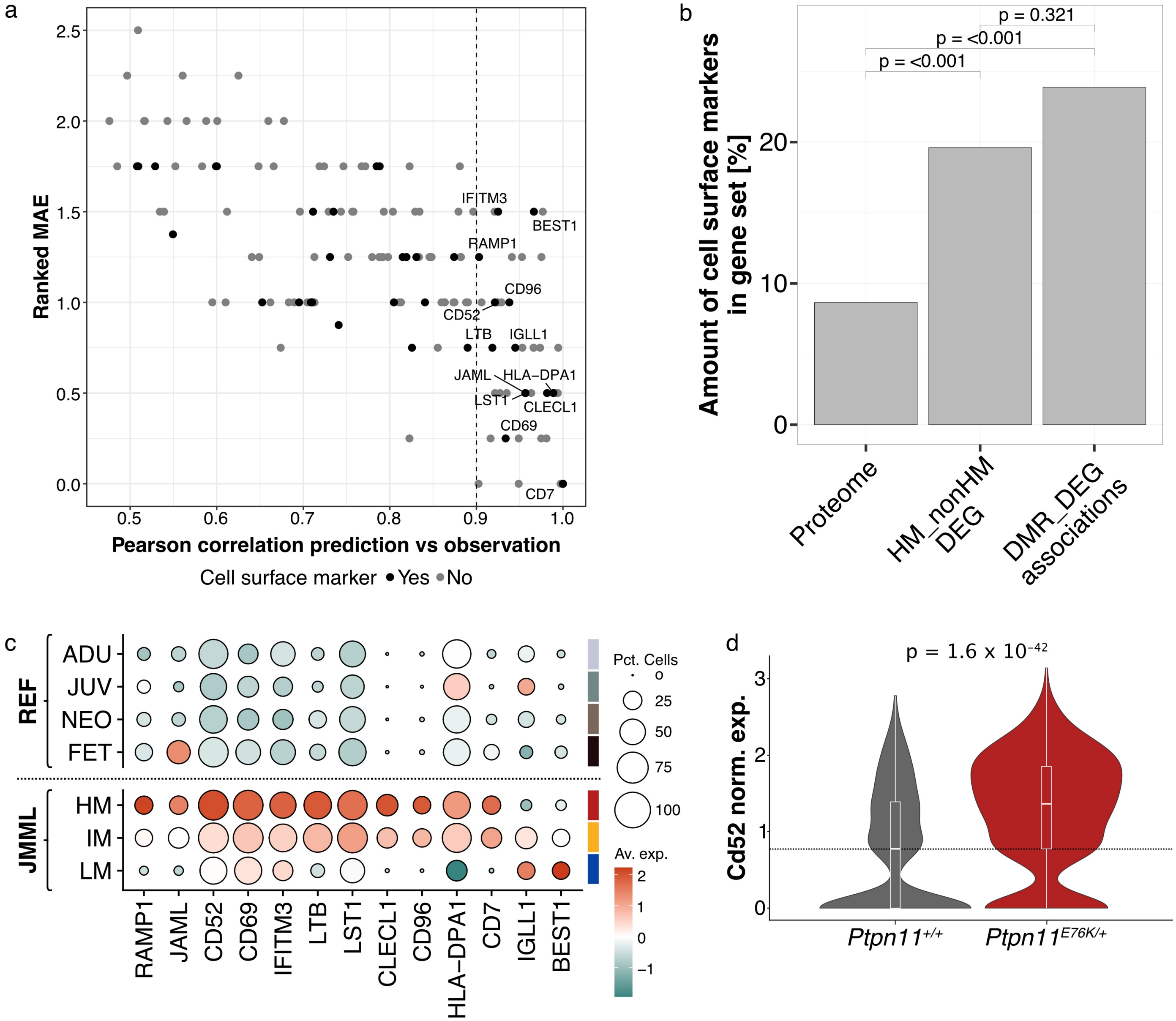

a
Ranked MAE
Pearson correlation prediction vs observation
Cell surface marker • Yes • No
IFITM3
BEST1
RAMP1
CD96
CD52
LTB
IGLL1
JAML
HLA−DPA1
LST1
CLECL1
CD69
CD7
b
Amount of cell surface markers in gene set [%]
p = 0.321
p = <0.001
p = <0.001
Proteome
HM_nonHM DEG
DMR_DEG associations
c
REF
ADU
JUV
NEO
FET
JMML
HM
IM
LM
RAMP1
JAML
CD52
CD69
IFITM3
LTB
LST1
CLECL1
CD96
HLA−DPA1
CD7
IGLL1
BEST1
Pct. Cells
Av. exp.
d
p = 1.6 x 10−42
Cd52 norm. exp.
Ptpn11+/+
Ptpn11E76K/+

**Extended Data Figure 11**: **Integrative analysis of methylation and transcription changes in JMML HSCs reveals novel prognostic biomarkers and therapeutic targets including CD52. a-b,** DMR-DEG associations to determine genes which subgroup-specific expression that can be explained by differential methylation. HM vs non-HM DMRs were associated with HM vs non-HM DEGs applying a generalized linear model with stepwise feature elimination using AIC. Per DEG, DMRs in a region 100 kb up- and downstream of the TSS were considered for potential associations. **a,** GLM results including all genes of which expression changes are associated with methylation changes across subgroups. Lower ranked MAE values represent genes that better fit the model. Pearson correlation between prediction and observation was used to calculate the confidence of the DMR-DEG associations. All genes above 0.9 were considered as candidates for further analysis. **b,** Amount of surface markers across gene sets reveals an enrichment of differentially expressed surface markers in JMML stem cells. As a reference, the previously published in silico human surfaceome was used[14]. 'DEG' are all HM vs. nonHM DEGs (Figure 2.I), 'DMR_DEG' are genes that were identified in glm analysis. **c,** Dot plot summarizing the scaled normalized expression of all DMR-associated high-confidence candidate surface marker genes in JMML and healthy reference HSCs. Circle size represents the percentage of cells expressing the gene. **d,** Violin plot of normalized single-cell RNA-seq expression of Cd52 in murine HSCs of *Ptpn11*$^{+/+}$ and *Ptpn11*$^{E76K}$ mice.

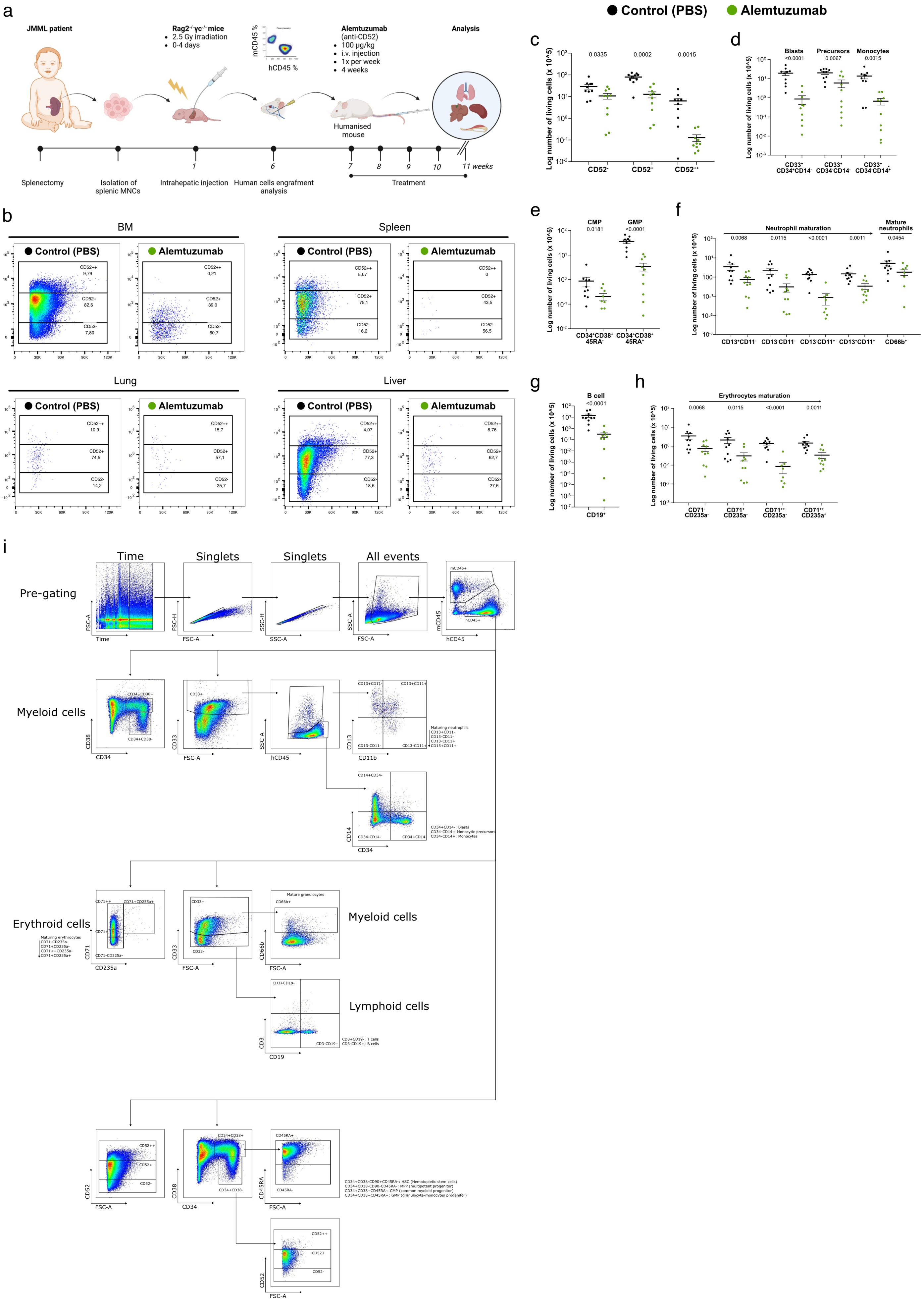
Control (PBS)
Alemtuzumab
a
JMML patient
Rag2-/-γc-/- mice
2.5 Gy irradiation
0-4 days
mCD45 %
hCD45 %
Alemtuzumab
(anti-CD52)
100 µg/kg
i.v. injection
1x per week
4 weeks
Analysis
Humanised mouse
Splenectomy
Isolation of splenic MNCs
Intrahepatic injection
Human cells engrafment analysis
Treatment
11 weeks
b
BM
Spleen
Lung
Liver
c
d
Blasts
Precursors
Monocytes
e
CMP
GMP
f
Neutrophil maturation
Mature neutrophils
g
B cell
h
Erythrocytes maturation
Log number of living cells (x 10^5)
i
Pre-gating
Time
Singlets
All events
Myeloid cells
Maturing neutrophils
CD34+CD14-: Blasts
CD34-CD14-: Monocytic precursors
CD34-CD14+: Monocytes
Erythroid cells
Maturing erythrocytes
Mature granulocytes
Lymphoid cells
CD3+CD19-: T cells
CD3-CD19+: B cells
CD34+CD38-CD90+CD45RA-: HSC (Hematopietic stem cells)
CD34+CD38-CD90-CD45RA-: MPP (multipotent progenitor)
CD34+CD38+CD45RA-: CMP (common myeloid progenitor)
CD34+CD38+CD45RA+: GMP (granulocyte-monocytes progenitor)

**Extended Data Figure 12**: **Anti-CD52 (Alemtuzumab) treatment depletes total hematopoiesis and disrupts disease-propagation in JMML patient-derived xenografts (PDX). a,** Schematic of anti-CD52 (Alemtuzumab) treatment of JMML PDX. Created with BioRender.com **b,** FACS plots showing CD52 expression in different organs in anti-CD52 treated (alemtuzumab, green) and untreated (Control, black) mice. **c-h,** Logarithmic scale of absolute cell number per pair of femora of human $CD45^+$ cells in treated vs. untreated mice. **c,** Total human $CD45^+$ cells stratified by CD52 expression. **d-h,** Different hematopoietic lineages based on classical cell type surface marker expression: **(d)** monocytes and blasts; **(e)** myeloid precursors; **(f)** neutrophils; **(g)** B cells; **(h)** erythrocytes. **i,** Gating scheme for immunophenotypic characterization of PDX mice.
